# Inputs to the locus coeruleus from the periaqueductal gray and rostroventral medulla shape opioid-mediated descending pain modulation

**DOI:** 10.1101/2023.10.10.561768

**Authors:** Susan T. Lubejko, Giulia Livrizzi, Janki Patel, Jean C. Yung, Tony L. Yaksh, Matthew R. Banghart

## Abstract

The supraspinal descending pain modulatory system (DPMS) shapes pain perception via monoaminergic modulation of sensory information in the spinal cord. However, the role and synaptic mechanisms of descending noradrenergic signaling remain unclear. Here, we establish that noradrenergic neurons of the locus coeruleus (LC) are essential for supraspinal opioid antinociception. Unexpectedly, given prior emphasis on descending serotonergic pathways, we find that opioid antinociception is primarily driven by excitatory output from the ventrolateral periaqueductal gray (vlPAG) to the LC. Furthermore, we identify a previously unknown opioid-sensitive inhibitory input from the rostroventromedial medulla (RVM), the suppression of which disinhibits LC neurons to drive spinal noradrenergic antinociception. We also report the presence of prominent bifurcating outputs from the vlPAG to the LC and the RVM. Our findings significantly revise current models of the DPMS and establish a novel supraspinal antinociceptive pathway that may contribute to multiple forms of descending pain modulation.

**Teaser:** Convergent synaptic activation of noradrenergic neurons in the locus coeruleus drives systemic opioid antinociception.

## Introduction

The midbrain and brainstem descending pain modulatory system (DPMS) alters spinal outflow of ascending nociceptive signals. Current models of the DPMS emphasize excitatory projections from ventrolateral periaqueductal gray (vlPAG) to the rostroventromedial medulla (RVM), which sends projections to the spinal cord dorsal horn that bidirectionally modulate incoming noxious sensory information through serotonergic and opioidergic mechanisms (*1*, *2*). Endogenous and exogenous opioids are thought to produce analgesia via disinhibition of vlPAG-to-RVM projection neurons (*3–5*). The RVM and, to some extent, the PAG, are comprised of On-, Off-, and neutral cells, which are defined electrophysiologically by their activity preceding a nocifensive response (*6–8*).

In addition to serotonin, rodent intrathecal pharmacological studies implicate spinal noradrenaline (NA) signaling in DPMS-driven and opioid-mediated pain suppression (*9–13*). Spinal serotonin can both suppress and facilitate nociceptive processing and pain behaviors (*14–16*), whereas spinal NA is more consistently reported as antinociceptive. Previous work implicates midbrain and brainstem catecholaminergic cells groups as the likely sources of spinal NA, especially the locus coeruleus (LC), also known as the A6 nucleus, which sends both ascending projections from dorsal LC to supraspinal areas, and descending projections from ventral LC to the spinal cord (*17–24*). Stimulation of ventral LC in rats causes antinociception that depends on spinal NA, while stimulation of dorsal LC is pronociceptive or anxiety-producing (*17*, *25*, *26*). Yet the role of LC in systemic opioid antinociception remains an open question, as the literature on this topic is sparse and conflicting (*27*, *28*).

Anatomical studies have identified projections from the PAG to the LC (*29*). Surprisingly, the synaptic features of these projections and their relevance to pain modulatory behavior have not been established. Although a rabies tracing study reported inputs to LC-NA neurons from vlPAG, and a slice electrophysiology study found an excitatory connection from lateral PAG to LC-NA neurons (*30–32*), *in vivo* electrophysiological recordings reported only weak, sparse input that predominantly *inhibits* LC discharge (*33*). Furthermore, vlPAG-to-LC neurons were reported to be inhibited by mu opioids (*31*), which conflicts with the notion that vlPAG output can drive descending LC neurons in the context of opioid analgesia. Thus, whether vlPAG contributes to the activation of LC to support opioid antinociception remains entirely unclear.

An additional complexity relates to the potential role of the RVM in influencing pain processing via synaptic communication with the LC. One anatomical study reports projections from both the raphe magnus and paragigantocellular nucleus, both components of the medial RVM, to the LC and pericoerulear region (*34*), but the sign and functional significance of this anatomical pathway remains unexplored. There is a strong body of literature demonstrating antinociceptive input to LC from excitatory neurons in the rostral ventro*lateral* medulla (RVLM) (*36*, *37*) and more recently, noradrenergic excitatory neurons in the caudal ventrolateral medulla (cVLM) (*38*). Yet, to our knowledge, there is no information regarding the nature and behavioral relevance of medial RVM inputs to the LC.

In this study, we use virally– and genetically-mediated anatomical, electrophysiological, and behavioral methods in mice to establish the circuit elements by which the DPMS recruits the LC to produce opioid antinociception. Our findings reveal the necessity of vlPAG input to the LC for systemic morphine antinociception and uncover an unexpected inhibitory input from the RVM that impacts nociception. These findings place new emphasis on noradrenergic signaling from the LC in opioid analgesia and reveal a novel antinociceptive pathway within the DPMS.

## Results

### Spinal NA release from the LC underlies systemic morphine antinociception

Intrathecal pharmacological experiments in rats point to a role for spinal noradrenergic and serotonergic signaling in systemic morphine antinociception (*13*). We first set out to confirm that spinal NA contributes to systemic morphine antinociception in mice by administering multiple doses of systemic morphine (5, 12, and 20 mg/kg, *s.c.*) in conjunction with intrathecal injections of the vehicle saline, the alpha-adrenergic antagonist phentolamine (5 μg), or, for comparison, the opioid antagonist naltrexone (5 μg) (**Figure 1A**). Systemic morphine with intrathecal saline administration increased hot plate withdrawal latencies in a dose-dependent manner (**Figure 1B**). At all morphine doses, intrathecal phentolamine blunted the resulting antinociception, and intrathecal naltrexone reduced it to an even greater degree (**Figure 1B-C**). These results indicate that spinal NA plays a critical role in the expression of systemic morphine antinociception, especially at low doses of morphine. In contrast, spinal opioid signaling, whether via direct spinal actions of morphine or DPMS-driven endogenous opioid release, is required for systemic morphine antinociception at all doses.

**Figure 1.**
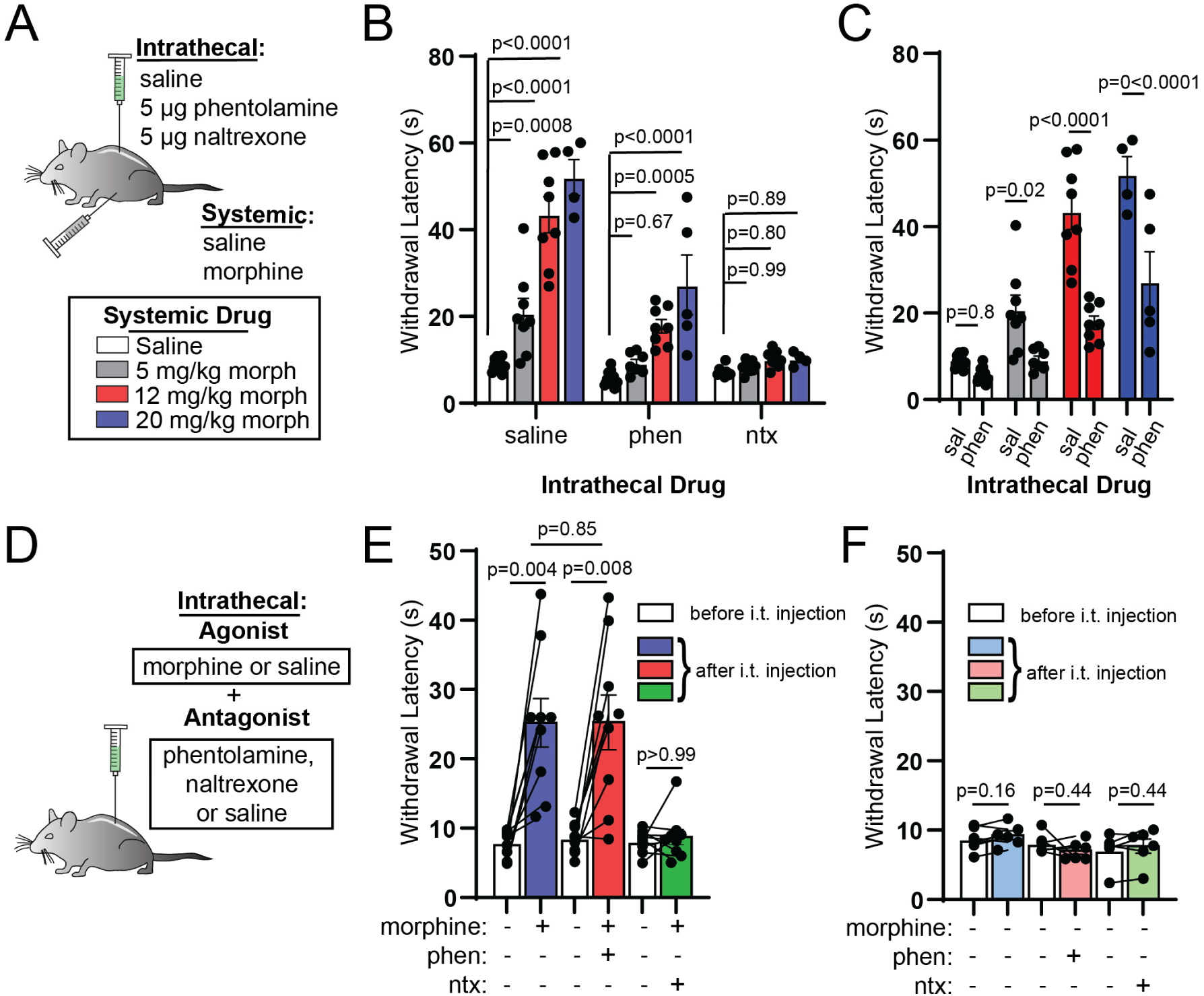
Intrathecal noradrenergic and opioidergic antagonists attenuate systemic morphine antinociception. **A**. Schematic of intrathecal and systemic injection combinations and morphine doses. **B.** Hot plate (52°C) withdrawal latencies resulting from increasing doses (0, 5, 12, 20 mg/kg) of subcutaneous morphine grouped by intrathecal antagonist administered (saline vs. 5 μg phentolamine vs. 5 μg naltrexone; ordinary two-way ANOVA with post hoc Tukey’s multiple comparisons test; intrathecal drug effect, p<0.0001, F(2,81)=95.35; morphine dose effect, p<0.0001, F(3,81)=56.61; interaction, p<0.0001, F(6,81)=16.75). **C.** Same data as in B from intrathecal saline and phentolamine groups organized by systemic morphine dose to facilitate comparisons at each dose (ordinary two-way ANOVA with post hoc Sidak’s multiple comparisons test; intrathecal drug effect, p<0.0001, F(1,52)=63.07; morphine dose effect, p<0.0001, F(3,52)=50.55; interaction, p=0.0002, F(3,52)=8.037). **D.** Schematic of intrathecal coadministration of morphine (2.5 μg) or saline with saline, phentolamine (5 μg) or naltrexone (5 μg). **E.** Hot plate withdrawal latencies before (white) and after (blue: saline, red: phentolamine, green: naltrexone) intrathecal injection of morphine and antagonist (pre– vs. post-intrathecal injection, n=9 pairs each, two-sided Wilcoxon matched-pairs signed rank test; post-intrathecal morphine + saline vs. post-intrathecal morphine + phentolamine, n=9 saline, n=9 phentolamine, two-sided Mann-Whitney test). **F.** Hot plate withdrawal latencies before (white) and after (light blue: saline, pink: phentolamine, light green: naltrexone) intrathecal injection of saline and antagonist (pre– vs. post-intrathecal injection, n=6 pairs each, two-sided Wilcoxon matched-pairs signed rank test). All bar graphs depict mean ± SEM and include individual experimental replicate values.

Direct spinal administration of morphine produces antinociception(*39*). It is therefore possible that morphine acts within the spinal cord to stimulate NA release, either from descending noradrenergic terminals or from unidentified local NA neurons. To determine whether morphine-driven spinal NA signaling originates from a spinal or supraspinal source, we intrathecally administered morphine (2.5 μg) along with saline, phentolamine, or naltrexone, prior to the hot plate assay (**Figure 1D**). Consistent with a supraspinal source of spinal NA, we found that the effect of intrathecal morphine was not blocked by intrathecal phentolamine, whereas intrathecal naltrexone blocked the antinociception (**Figure 1E**). In the absence of morphine, these antagonists did not significantly change withdrawal latency (**Figure 1F**).

Previous studies have identified several supraspinal noradrenergic structures that project to the dorsal horn of the spinal cord (*18*, *20–24*). Whereas the A5 and A7 nuclei contain only a small fraction of dorsal horn-projecting NA neurons, the LC is the primary source of spinal NA (*18*). Stimulation of the spinally-projecting ventral subdivision of LC produces spinal NA-dependent antinociception in mice and rats (*17*, *25*, *26*). To determine whether systemic morphine increases the activity of LC-NA neurons, we measured expression of the immediate early gene c-Fos in response to subcutaneous morphine (10 mg/kg) or saline treatment in *DBH-cre:tdTom* mice that express the fluorescent protein tdTomato in dopamine beta-hydroxylase (DBH)-positive neurons. Consistent with LC activation, systemic morphine caused a significant increase in c-Fos expression in LC neurons (**Figure 2A-B**), which was significantly blocked by systemic opioid receptor antagonist naloxone (10 mg/kg, s.c.) pretreatment. This presents a paradox, as LC neurons express the mu opioid receptor (MOR), which typically inhibits neuronal output in receptor-expressing neurons. A possible explanation is that the c-Fos induction actually results from direct engagement of mitogen-activated protein kinase cascades downstream of MOR activation (*40*). However, morphine-induced c-Fos expression was unchanged in *DBH-cre::Oprm1^fl/fl^* mice (*41*), which lack MOR in NA neurons (**Figure S1A-B**). Taken together, these data suggest that while MOR expression in LC neurons themselves is not responsible for morphine-induced c-Fos expression, opioid receptors present in the circuit upstream from these neurons are responsible for LC c-Fos expression, possibly via disinhibition of LC activity.

**Figure 2.**
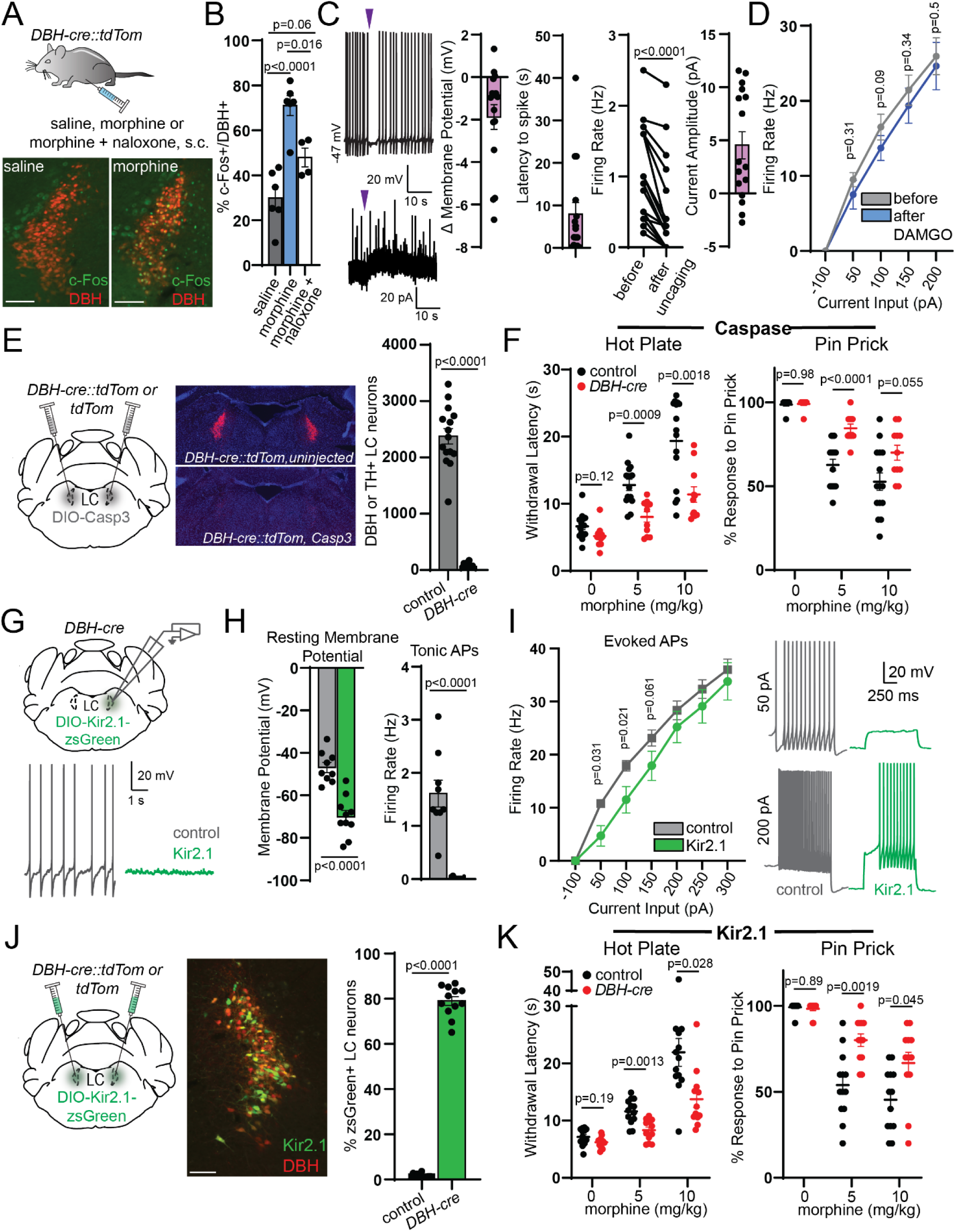
LC activity is required for systemic morphine antinociception. **A**. Subcutaneous saline, 10 mg/kg morphine, and 10 mg/kg morphine + 10 mg/kg naloxone injections and representative images of resulting c-Fos (green) immunohistochemistry in *DBH-cre::tdTom* mice. Scale bars 150 μm. **B.** Percentage of DBH-positive LC neurons that colocalize with green c-Fos signal (5-8 LC images analyzed per mouse; n=6 saline, n=6 morphine, n=4 morphine + naloxone; saline = 29.9 ± 5.1%, morphine = 71.0 ± 4.5%, morphine + naloxone = 47.9 ± 4.2%; Ordinary One-way ANOVA with post hoc Tukey’s multiple comparisons test, p<0.0001, F(2,13) = 20.97)). **C.** Left: representative traces of CNV-Y-DAMGO light-evoked uncaging (purple arrows: 50 ms, 365 nm LED, 84 mW) during current clamp (top) and voltage clamp (bottom) recordings in whole cell patch clamp from LC neurons in wild type mice. Right: LC neuron opioid response characterized by change in membrane potential after light flash, pause in firing calculated as latency to first spike after light flash, tonic firing rate during the 10 seconds before and 10 seconds after light flash (p<0.0001, two-sided Wilcoxon matched-pairs signed rank test), and evoked outward current amplitude (n=16 cells each). **D.** f-I curves recorded from LC neurons in wild type mice before and after bath application of 1 μM DAMGO (n=7 cells; two-sided Wilcoxon matched-pairs signed rank test at 50 pA, 100 pA, 150 pA, and 200 pA). **E.** Injection schematic, representative images and quantification of neuronal ablation of mice after bilateral injection of AAV-DIO-Casp3 in LC (n=11 *DBHcre:TdTom*, n=15 *TdTom* control; *DBH-cre* = 79 ± 15 cells, control = 2376 ± 137 cells; two-sided Mann-Whitney test). **F.** Left, hot plate paw withdrawal latencies of control and *DBH-cre* mice following subcutaneous administration of 0, 5, and 10 mg/kg morphine (Two-way repeated measures ANOVA with post hoc Sidak’s multiple comparisons test; n=11 *DBHcre:TdTom* n=15 *TdTom* control; morphine dose effect, p<0.0001, F(1.365,32.75)= 78.80, genotype effect, p=0.0006, F(1,24)=15.67, morphine dose x genotype interaction, p=0.0004, F(2,48)=9.328). Right, Response to 10 hind paw pin pricks following morphine administration (Two-way repeated measures ANOVA with post hoc Sidak’s multiple comparisons test; n=11 *DBHcre:TdTom* n=15 *TdTom* control; morphine dose effect, p<0.0001, F(1.690,40.56)=72.20, genotype effect, p=0.0006, F(1,24)=15.52, morphine dose x genotype interaction, p=0.0037, F(2,48)=6.294). **G.** Top, Slice electrophysiology recordings from Kir2.1-positive and –negative LC neurons. Bottom, representative traces of tonic AP firing in control vs. Kir2.1-expressing neurons. **H.** Left, resting membrane potential of control vs. Kir2.1-expressing neurons (n=10 Kir2.1-positive, n=9 control; Kir2.1 RMP = –70.1 ± 3.0 mV, control RMP = –46.9 ± 2.4mV, two-sided Mann-Whitney test). Right, tonic AP firing rate of control vs. Kir2.1-expressing neurons (Kir2.1 tonic firing rate = 0.006 ± 0.006 Hz, control tonic firing rate = 1.61 ± 0.25 Hz, two-sided Mann-Whitney test). **I.** Left, f-I curves of control vs. Kir2.1-expressing neurons for 7 current steps (n=10 Kir2.1-positive, n=9 control; two-sided Mann-Whitney test). Right, representative traces of evoked AP firing from a control (grey) and a Kir2.1-expressing (green) neuron at 50 pA (top) and 200 pA (bottom) current steps. **J.** Injection schematic, representative image of zsGreen expression in the LC of a *DBH-cre::tdTom* mouse (scale bar = 150 μm) and quantification of viral coverage given as % tdTom LC neurons labeled by zsGreen (n=12 *DBHcre:TdTom*, n=13 *TdTom* control; control = 1.2 ± 0.3%, *DBH-cre* = 79.0 ± 2.0%; two-sided Mann-Whitney test). **K.** Left, hot plate paw withdrawal latencies of control and *DBH-cre* mice following 0, 5, and 10 mg/kg morphine, s.c. (Two-way repeated measures ANOVA with post hoc Sidak’s multiple comparisons test; n= 12 *DBHcre:TdTom*, n=13 *TdTom* control; hot plate: morphine dose effect, p<0.0001, F(1.092,25.11)=46.27, genotype effect, p=0.001, F(1,23)=14.30, morphine dose x genotype interaction, p=0.013, F(2,46)=4.790). Right, Response to 10 hind paw pin pricks following morphine (Two-way repeated measures ANOVA with post hoc Sidak’s multiple comparisons test; n= 12 *DBHcre:TdTom*, n=13 *TdTom* control; morphine dose effect, p<0.0001, F(1.695,38.99)=58.23, genotype effect, p=0.0004, F(1,23)=17.20, morphine dose x genotype interaction, p=0.0044, F(2,46)=6.131). Data in each graph reported as mean ± SEM.

In rat brain slices, MOR agonists strongly hyperpolarize LC neurons via activation of G protein-coupled inward rectifier K^+^ (GIRK) channels (*42*, *43*). Because nearly all prior observations of MOR-mediated dampening of LC neuronal excitability were made in rats, we asked if mouse LC-NA neurons are similarly affected. Using fluorescent *in situ* hybridization, we first confirmed that *Oprm1* and *TH* transcripts, which encode the MOR and tyrosine hydroxylase, respectively, colocalize in the LC of C57Bl6/J mice (**Figure S1D-E**). We next investigated the effects of the MOR agonist DAMGO on LC neuronal excitability using whole cell electrophysiological recordings in acute mouse brain slices. Bath applied DAMGO requires minutes to equilibrate in slices, which can obscure subtle effects on membrane properties. We therefore used the photocaged DAMGO derivative CNV-Y-DAMGO (1 μM) to produce time-locked DAMGO concentration jumps in response to millisecond flashes of light(*44*, *45*). In current clamp recordings, despite the use of a strong optical stimulus, CNV-Y-DAMGO photoactivation generated only small hyperpolarizations that were accompanied by brief pauses in spontaneous firing and minor subsequent reductions in firing rate (**Figure 2C**). In voltage clamp recordings, DAMGO photorelease evoked small outward currents (**Figure 2C**). Consistent with these modest effects, bath application of DAMGO produced no change in evoked action potential firing (**Figure 2D**). These small effects stand in stark contrast to the several-hundred pA currents and deep hyperpolarizations evoked by MOR agonists in rat LC, and are consistent with the smaller GIRK currents previously observed in mice on a different genetic background (*46*).

These results suggest that direct MOR-mediated inhibition of LC firing may be insufficient to counteract synaptic drive of LC neurons by upstream circuits in mice. Consistent with this notion, genetic removal of MORs from LC-NA neurons did not alter systemic morphine antinociception on the hot plate (**Figure S1C**).

To determine if the LC contributes to morphine antinociception, we employed two loss-of-function approaches. First, we specifically ablated LC-NA neurons via bilateral injection of cre-dependent Caspase 3 (Casp3) into the LC of either *DBH-cre:tdTom* mice or control *tdTom* littermates. Transduction with Casp3 led to near-complete ablation of LC-NA neurons (**Figure 2E**). In addition to assessing thermal nociception, we used a pin prick assay to measure mechanical nociception. Somewhat surprisingly, LC ablation had no effect on baseline nocifensive responses in the absence of morphine. However, in comparison to control mice, the morphine-induced increase in hot plate withdrawal latency was strongly attenuated in *DBH-cre:tdTom* mice at both doses of morphine (5 and 10 mg/kg). In addition, the morphine-induced decrease in pin prick response was significantly attenuated at 5 mg/kg morphine and trended toward significance at 10 mg/kg (**Figure 2F**).

Casp3-mediated cell ablation is a severe manipulation that may lead to neuroinflammation and/or a loss of structural integrity. We therefore sought to suppress LC activity by overexpressing Kir2.1, a constitutively active inwardly-rectifying K^+^ channel, which hyperpolarizes neurons and decreases their excitability. We first analyzed the effect of Kir2.1 overexpression on LC neuronal excitability using brain slice electrophysiology 3 weeks after unilateral LC injection of AAV-DIO-Kir2.1-zsGreen in *DBH-cre* mice. Kir2.1 overexpression decreased resting membrane potential and abolished tonic action potential firing compared to anatomically-identified LC neurons in the uninjected hemisphere (**Figure 2G-H**). We also observed a decrease in evoked action potential firing in response to injection of 50 and 100 pA current steps (**Figure 2I**). Taken together, Kir2.1 overexpression effectively decreases both tonic and evoked action potential firing in LC neurons.

For behavioral testing, we injected *DBH-cre:tdTom* or *tdTom* mice bilaterally in the LC with AAV-DIO-Kir2.1-zsGreen. Histological analysis revealed high transduction of LC neurons in cre-expressing mice and negligible expression in cre-negative littermates (**Figure 2J**). Kir2.1 overexpression suppressed morphine antinociception on both the hot plate and pin prick assays at both doses of morphine (**Figure 2K**). Notably, neither Casp3 nor Kir2.1 produced a difference in locomotion or basal anxiety in the elevated plus maze (**Figure S2A-B**). Overall, these results support a critical role for the LC in systemic morphine antinociception.

### vlPAG gates LC-mediated morphine antinociception

While LC is crucial for systemic morphine antinociception, it remains unclear how LC activity is recruited by upstream structures in response to opioid drugs. Classic models of opioid antinociception involve the disinhibition of vlPAG output neurons via activation of MORs expressed on inhibitory terminals and/or local GABA neurons within the vlPAG (*3*, *5*).

Consistent with this, the antinociception produced by local morphine infusion into rat PAG partially depends on spinal NA (*12*). Furthermore, several PAG subregions, including the vlPAG, send excitatory projections to the LC and pericoerulear region (*29–32*). Although optogenetic and chemogenetic activation of vlPAG glutamatergic neurons produces antinociception in mice (*47*, *48*), whether this is mediated by the LC, and whether vlPAG glutamatergic neurons support morphine antinociception, is not known.

If vlPAG is an important upstream mediator of morphine antinociception, we reasoned that inhibiting vlPAG output should attenuate the antinociception produced by 5 mg/kg morphine, which we found to rely heavily on the LC. To probe the role of vlPAG in morphine antinociception, we bilaterally expressed the inhibitory DREADD HM4D (*49*) in vlPAG*^VGLUT2-cre^*neurons and inhibited their output with systemic CNO (3 mg/kg, i.p.) in the absence and presence of morphine (5 mg/kg, *s.c.*) (**Figure 3A**). In untransduced control mice, CNO had no effect on morphine antinociception. In contrast, CNO completely prevented morphine antinociception in HM4D-expressing mice (**Figure 3B**).

**Figure 3.**
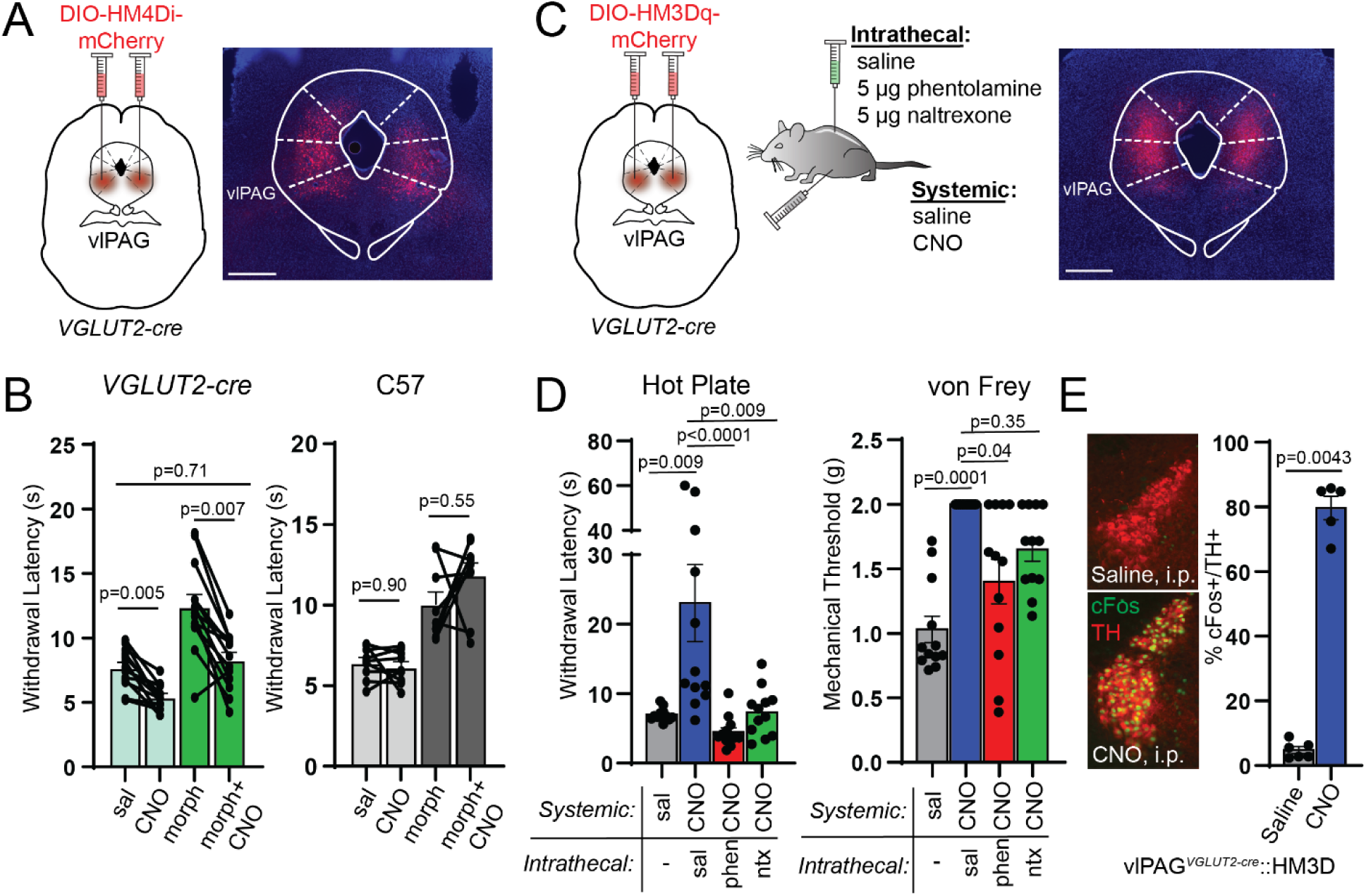
vlPAG activity is required for systemic morphine antinociception and drives spinal NA-dependent antinociception. **A**. Bilateral AAV-DIO-HM4Di-mCherry injections in vlPAG of *VGLUT2-cre* mice with representative image of viral expression. Scale bar 500 μm. **B.** Left, hot plate withdrawal latencies of vlPAG*^VGLUT2-cre^*::HM4D mice administered 3 mg/kg CNO, i.p. vs. saline without (light green bars) and with 5 mg/kg morphine, s.c. (dark green bars) (Two-way repeated measures ANOVA with post hoc Tukey’s multiple comparisons test; n=13 mice; sal vs. CNO effect, p<0.0001, F(1,12)=56.93, morphine effect, p<0.0001, F(1,12)=43.25, morphine x CNO interaction, p=0.048, F(1,12)=4.837). Right, hot plate withdrawal latencies of non-virus injected C57 controls (Two-way repeated measures ANOVA with post hoc Tukey’s multiple comparisons test; n=9 mice; sal vs. CNO effect, p=0.32, F(1,8)=1.143, morphine effect, p<0.0001, F(1,8)=75.02). **C.** Bilateral AAV-DIO-HM3Dq-mCherry injections in vlPAG of *VGLUT2-cre* mice and combinations of systemic CNO (3 mg/kg, i.p.) with intrathecal antagonists and representative image of viral expression. Scale bar 500 μm. **D.** Left, hot plate withdrawal latencies after systemic saline or CNO and intrathecal saline, phentolamine (5 μg), or naltrexone (5 μg) (Friedman test with post hoc Dunn’s multiple comparisons test, n=12 subjects, p<0.0001, Friedman statistic = 26.40). Right, von Frey mechanical thresholds (Friedman test with post hoc Dunn’s multiple comparisons test, n=12 subjects, p=0.0001, Friedman statistic = 21.00). **E.** Representative images of c-Fos immunohistochemistry in TH-positive LC neurons of vlPAG*^VGLUT2-cre^*::HM3D mice after systemic injection of saline (top) or CNO (bottom). Right, % of TH-positive LC neurons that colocalize with green c-Fos signal (5-8 images analyzed per mouse; n= 6 saline, n=5 CNO; saline = 4.8 ± 1.1%, CNO = 79.6 ± 3.6%, two-sided Mann-Whitney test). Data in each graph reported as mean ± SEM.

To determine if spinal NA signaling mediates vlPAG-driven antinociception, we bilaterally activated vlPAG*^VGLUT2-cre^* neurons with the excitatory DREADD HM3D in conjunction with intrathecal NA antagonism (**Figure 3C**). Consistent with prior work (*48*), systemic CNO (3 mg/kg, i.p.) increased hot plate withdrawal latencies and von Frey mechanical thresholds.

Strikingly, this effect was either completely or partially abolished on the thermal and mechanical assays, respectively, by intrathecal administration of either phentolamine or naltrexone (**Figure 3D**). However, we also observed that chemogenetic activation of vlPAG*^VGLUT2-cre^*neurons produced a state of active quiescence, in which mice exhibited a marked reduction in voluntary locomotion, distinct from freezing behavior (**Figure S3A**). This raises a concern that the apparent antinociception is simply a consequence of general locomotor suppression. However, the active quiescence persisted after intrathecal phentolamine and naltrexone administration, despite significant attenuation of the antinociception. Therefore, DPMS-driven descending antinociception and locomotor suppression are dissociable. Consistent with vlPAG driving the LC to produce spinal NA-dependent antinociception, we also found that chemogenetic activation of vlPAG*^VGLUT2-cre^* neurons increased c-Fos expression in TH-positive LC neurons (**Figure 3E**), yet CNO had no effect in the LC of untransduced wildtype mice (**Figure S3B**). Together, these results demonstrate that, like systemic morphine, vlPAG-driven antinociception requires spinal NA signaling. They also establish a role for spinal endogenous opioid signaling in vlPAG-driven antinociception, a point debated in previous literature (*9*, *50*).

### Anatomy of DPMS input to the LC

To uncover the circuit elements by which the DPMS may recruit the LC, we first mapped projections from the vlPAG and RVM to the LC by virally expressing tdTomato in neurochemically-defined cell types. To label glutamatergic neurons, we used *VGLUT2-cre* mice, as VGLUT2 is the primary vesicular glutamate transporter isoform in both structures. To label inhibitory axons, we used *VGAT-cre* mice, which express cre in both GABAergic and glycinergic neurons (*51*).

After unilateral injection of AAV-DIO-tdTom into the vlPAG of either *VGLUT2-cre* or *VGAT-cre* mice, we observed sparse fibers in the LC somatic region and strong labeling in the surrounding pericoerulear region, including a medial zone corresponding to Barrington’s nucleus(*52*, *53*) and a dorsolateral zone near the 4^th^ ventricle (**Figure 4A-E**). Quantification of pixel intensity of green TH+ LC neurons and red fibers supports dense vlPAG projections just medial to the LC (**Figure 4F-G**). Consistent with axo-dendritic inputs to LC-NA neurons occurring within the pericoerulear region and evidence of pericoerulear interneurons that modulate LC activity(*54–56*), these results establish anatomical substrates by which vlPAG could bidirectionally control LC activity.

**Figure 4.**
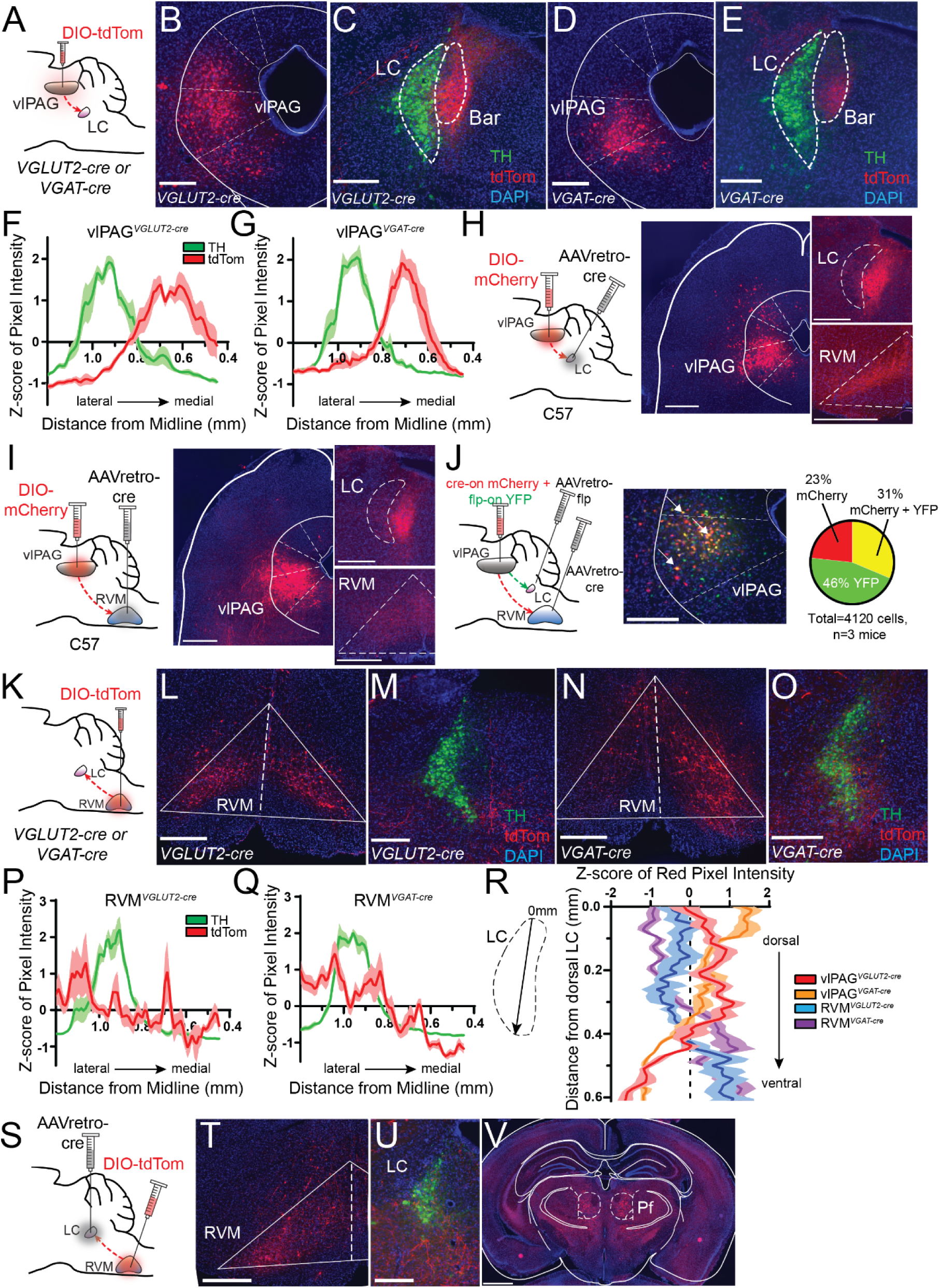
Anatomical characterization of inputs to LC from vlPAG and RVM. **A**. Injection of AAV-DIO-tdTom in vlPAG of *VGLUT2-* or *VGAT-cre* mice. **B.** Injection site in *VGLUT2-cre* mouse. **C.** vlPAG glutamatergic terminals (red) in pericoerulear region defined by TH immunohistochemistry (green). Delineations taken from the Paxinos & Franklin Brain Atlas show that the pericoerulear terminals are largely targeted to the medial Barrington’s nucleus. **D.** Injection site in *VGAT-cre* mouse. **E.** vlPAG GABAergic/glycinergic terminals in pericoerulear region. **B-E**. Scale bars 300 μm. **F.** Quantification of red (vlPAG glutamatergic terminals) and green (LC TH immunohistochemistry) pixel intensity in the LC and pericoerulear region normalized by z-score to account for fluorescence differences between animals and during imaging (n=6 LC slices from n=3 mice each). **G.** Same as F for vlPAG GABAergic/glycinergic terminals. **H.** Left: injections of AAVretro-cre in LC and AAV-DIO-mCherry in vlPAG of wild type mice to capture LC-projecting vlPAG neurons. Center: image of mCherry+ vlPAG neurons captured. Right: resulting terminals captured in LC and RVM. Scale bars 500 μm. **I.** Same as H for vlPAG neurons that project to RVM. Scale bars 500 μm. **J.** Left: Orthogonal recombinase strategy to label vlPAG neurons that project to RVM and LC. Center: representative image of mCherry (red) and YFP (green) viral labeling in vlPAG with neurons co-expressing both fluorophores appearing yellow. White arrows indicate examples of double labeled neurons. Scale bar 300 μm. Right: quantification of mCherry and YFP labeling in vlPAG. **K.** Injection of AAV-DIO-tdTom in RVM of *VGLUT2-* or *VGAT-cre* mice. **L.** Injection site in *VGLUT2-cre* mouse. **M.** RVM glutamatergic terminals (red) in LC and pericoerulear region defined by TH immunohistochemistry (green). **N.** Injection site in *VGAT-cre* mouse. **O.** RVM GABAergic/glycinergic terminals in LC and pericoerulear region. **L-O.** Scale bars 300 μm. **P.** Quantification of RVM glutamatergic terminals by pixel intensity z-score similar to F (n=6 LC slices from n=3 mice each). **Q.** Same as P for RVM GABAergic/glycinergic terminals. **R.** Quantification of red pixel intensity normalized by z-score across the dorsal to ventral axis of LC for all four projection origin and cell type combinations (n=6 LC slices from n=3 mice each). **S.** Injections of AAVretro-cre in LC and AAV-DIO-tdTom in RVM of wild type mice to capture LC-projecting RVM neurons. **T.** tdTom+ RVM neurons captured. **U.** Resulting terminals (red) captured in LC (green) and pericoerulear region. Scale bar 300 μm. **V.** tdTom+ fibers located in bilateral parafascicular nucleus of the thalamus. Scale bar 1mm.

As expected, tdTom expression in vlPAG*^VGLUT2-cre^* neurons also labeled axons in the RVM (**Figure S4A**). We also observed prominent fluorescent axons in the RVM upon tdTom expression in vlPAG*^VGAT-cre^* neurons. The existence of inhibitory vlPAG-to-RVM projections has been established in rats but refuted in mice (**Figure S4B**) (*57–60*). We verified this inhibitory projection using a retro-FISH approach involving retrobead injection into RVM prior to retrobead-labeled vlPAG cell type identification using fluorescence *in situ* hybridization (**Figure S4C**). This analysis revealed that although the vast majority of vlPAG-to-RVM projection neurons contain transcripts encoding glutamatergic (but not GABAergic) markers, ∼15% contain only GABAergic markers (**Figure S4D**). These results establish that vlPAG sends inhibitory projections to the RVM in mice.

A previous study in mice suggested the presence of two distinct, non-overlapping populations of vlPAG neurons that project to the LC and RVM (*31*). We wondered if vlPAG-to-LC neurons send branches to other brain areas. To address this question, we fluorescently labeled vlPAG neurons that project to the LC using the retrograde virus AAVretro-cre in wild-type mice (**Figure 4H**).

Consistent with our previous results, we observed mCherry-positive fibers in the medial pericoerulear region. However, we also observed prominent axon fibers in the RVM, with no other apparent midbrain and brainstem targets. To confirm this result, we labeled vlPAG-to-RVM neurons using the same approach and again observed fluorescent axons in the medial pericoerulear region (**Figure 4I**). We verified these findings using a dual-color double-retrograde approach by injecting AAVretro-cre into the RVM, AAVretro-FlpO into the LC, and a mixture of cre– and flp-dependent mCherry and YFP reporter viruses in vlPAG. Importantly, the reporter virus titers were optimized to eliminate recombinase-independent expression and recombinase cross-talk. We observed mCherry and YFP co-expression in 31% of fluorescently labeled neurons (**Figure 4J**), which is likely an underestimate due to incomplete viral uptake. Taken together, these experiments establish that a large number of vlPAG output neurons send bifurcating axons to both the LC and RVM. Importantly, these results imply that vlPAG recruits the LC and RVM in an inseparable, parallel manner.

We next asked if we could detect projections from the RVM to the LC, which were reported a single prior study (*34*). Upon injecting AAV-DIO-TdTom into the RVM of *VGLUT2-cre* or *VGAT-cre* mice, red fluorescent fibers were found in the LC and pericoerulear region in innervation patterns distinct from vlPAG axons (**Figure 4K-O**). Notably, pixel intensity analysis revealed that RVM*^VGLUT2-cre^*axons were sparse within the LC somatic region and in the medial pericoerulear region, did not innervate Barrington’s nucleus, and innervated a region just lateral to LC. In contrast, RVM*^VGAT-cre^* axons appeared to strongly innervate the LC somatic region and symmetrically spill over only slightly into the medial and lateral pericoerulear regions (**Figure 4P-Q**). Additionally, analysis of fibers from all four projection origin– and cell type combinations along the dorsal-ventral axis within the LC somatic region suggested that vlPAG largely targets dorsal LC whereas RVM targets ventral LC (**Figure 4R**).

To determine if RVM-to-LC neurons send branching axons to other structures, we injected AAVretro-cre in LC and AAV-DIO-tdTom in RVM (**Figure 4S**). We observed fluorescent fibers in several brain regions other than the LC. Most prominent was a dense projection to the parafascicular nucleus of the thalamus, which also receives input from LC (*61*) (**Figure 4T-V**). Notably, we did not find fluorescent axons in the spinal cord dorsal horn, indicating that RVM-LC projection neurons belong to a population distinct from the spinally-projecting RVM neurons that directly modulate incoming noxious sensory information. Taken together, these results suggest that the LC receives neurochemically-diverse synaptic inputs from multiple hindbrain nodes within the DPMS.

### Synaptic properties of vlPAG and RVM input to the LC

To gain insight into the functional significance of these pathways, we optogenetically stimulated vlPAG or RVM axons in LC during acute brain slice electrophysiological recording from LC-NA neurons. To achieve high expression of channelrhodopsin (ChR2) in all projection cell types, we injected wild-type mice in either the vlPAG or RVM with a combination of AAVDJ-Ef1a-mCherry-IRES-cre and AAVDJ-Ef1a-DIO-ChR2-mCherry. After 3-4 weeks of expression, we prepared coronal slices and obtained whole-cell recordings from morphologically-identified LC neurons. We targeted cells that were located in the ventral half of LC in order to bias our recordings towards the spinally-projecting population (*18*) and confirmed that each recorded neuron exhibited spontaneous tonic firing at ∼1.5 Hz.

We first determined the net effect of optogenetic axon stimulation on LC neuron firing using a 2-second blue light stimulus (25 Hz). Upon activating vlPAG inputs, we observed a firing rate increase in 80% of recorded neurons, whereas 20% showed no significant change. Across the population, this resulted in an overall increase in firing rate during the stimulus (**Figure 5A-B**). In contrast, upon activating RVM inputs to LC, we observed a firing rate increase in in only 10% of the recorded neurons, and a decrease in 60%, whereas 30% were not modulated. Across the population, this resulted in an overall decrease in firing rate (**Figure 5C,D**).

**Figure 5.**
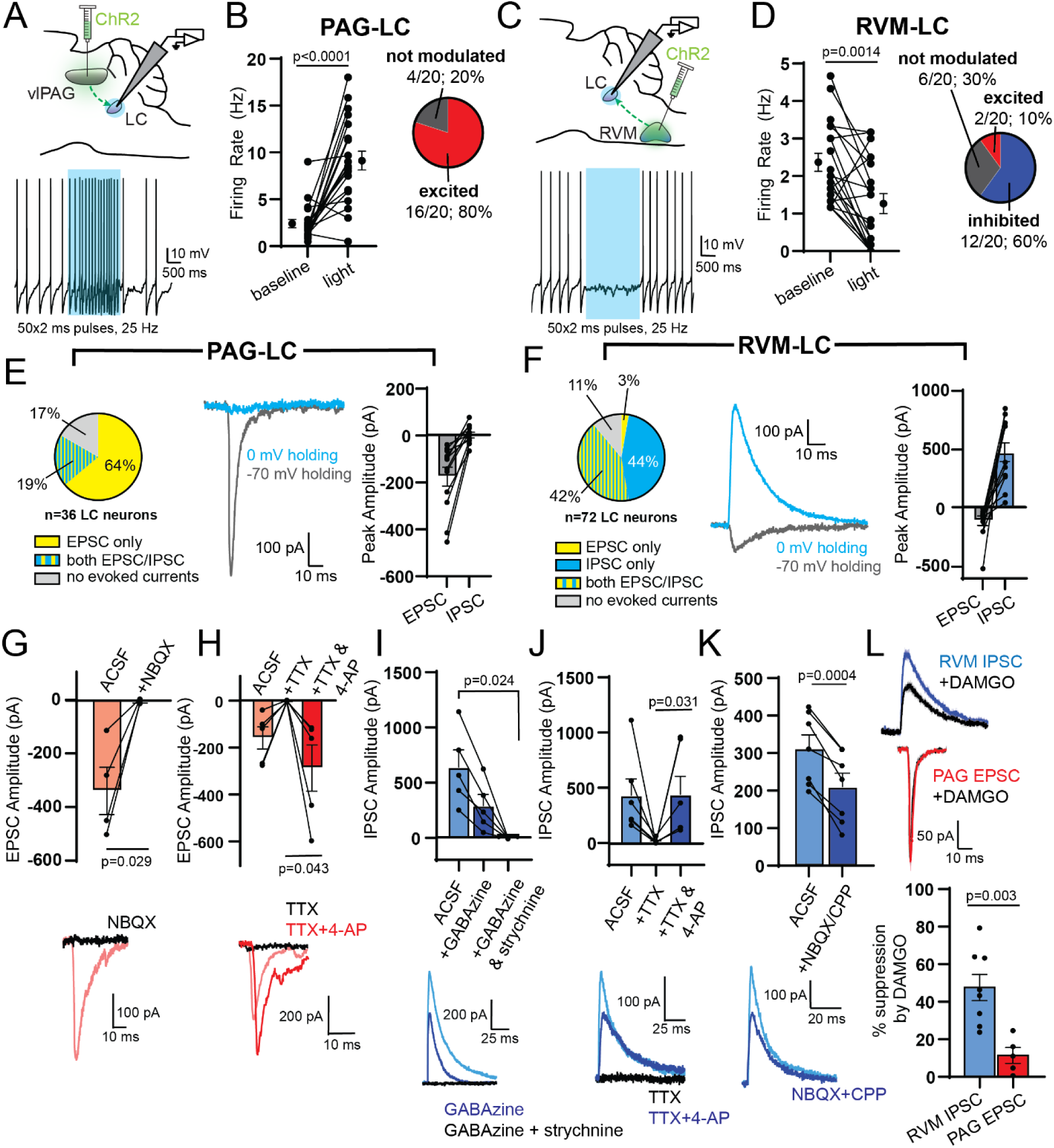
Electrophysiological characterization of inputs from vlPAG and RVM to LC. **A**. Top, viral injection of ChR2 into vlPAG for LC slice electrophysiology. Bottom, representative trace of tonic spiking before, during, and after a 2 second blue LED stimulus (470 nm, 50×2 ms pulses at 25 Hz, 18 mW). **B.** Left, firing rate before and during the light stimulus for 20 LC neurons (baseline = 2.4 ± 0.4 Hz, light on = 9.1 ± 1.0 Hz; two-sided Wilcoxon matched-pairs signed rank test). Right, recorded neurons were categorized as “excited” (a z-score of spiking during vs. before light >2), “inhibited” (a z-score of spiking during vs. before light < –2), or “not modulated.” **C.** Top, viral injection of ChR2 into RVM for LC slice electrophysiology. Bottom, representative trace of tonic spiking before, during, and after the blue LED stimulus. **D.** Left, firing rate before and during the blue light stimulus for 20 LC neurons (baseline = 2.3 ± 0.2 Hz, light on = 1.3 ± 0.3 Hz; two-sided Wilcoxon matched-pairs signed rank test). Right, categorization of recorded LC neurons as “excited”, “inhibited”, or “not modulated.” **E.** Left, proportions of LC neurons in which optically-evoked EPSCs, EPSCs and IPSCs or no evoked currents were present when stimulating vlPAG terminals (470nm, 1 x 5 ms pulse, 18 mW). Middle, representative example of oEPSC and oIPSC recorded in a single LC neuron via electrical isolation. Right, peak amplitude of oEPSCs and oIPSCs driven by vlPAG terminal stimulation (oEPSCs = –177.8 ± 40.6 pA, oIPSCs = –0.52 ± 12.7 pA, n=12 neurons). **F.** Left, proportions of LC neurons in which optically-evoked EPSCs, IPSCs, both EPSCs and IPSCs or no evoked currents were present when stimulating RVM terminals. Middle, representative example of oEPSC and oIPSC recorded in a single LC neuron via electrical isolation. Right, peak amplitude of oEPSCs and oIPSCs driven by RVM terminal stimulation (oEPSCs = –117.7 ± 39.9 pA, oIPSCs = 466.6 ± 79.5 pA, n=12 neurons). **G-K.** Top, summary bar graphs of oEPSC/IPSC amplitude. Bottom, representative examples. **G.** vlPAG-driven oEPSC amplitude before and after bath application of NBQX (10 μM) (ACSF = –337.1 ± 88.0 pA, +NBQX = –4.9 ± 4.4 pA; two-sided paired t-test: t=3.950, n=4 pairs). **H.** vlPAG-driven oEPSC amplitude before and after bath application of TTX (1 μM) and subsequent application of 4-AP (100 μM) (+TTX = –2.9 ± 1.5 pA, +TTX & 4-AP = –288.5 ± 98.4 pA; two-sided paired t-test: t=2.926, n=5 pairs). **I.** RVM-driven oIPSC amplitude before and after bath application of GABAzine (20 μM) and additional application of strychnine (10 μM) (ACSF = 636.3 ± 154.9 pA, +GABAzine = 286.6 ± 100.8 pA, +GABAzine & strychnine = –2.5 ± 1.7 pA; Repeated Measures One-way ANOVA with Dunnett’s multiple comparisons test, p=0.013, F(1.087,4.348)=16.32, n=5 cells). **J.** RVM-driven oIPSC amplitude before and after bath application of TTX (1 μM) and subsequent application of 4-AP (100 μM) (+TTX = 13.1 ± 9.8 pA, +TTX & 4-AP = 443.9 ± 164.6 pA; two-sided Wilcoxon matched-pairs signed rank test, n=6 pairs). **K.** RVM-driven oIPSC amplitude before and after bath application of NBQX (10 μM) + CPP (10 μM) (ACSF = 309.6 ± 36.1 pA, +NBQX/CPP = 207.6 ± 35.8 pA; two-sided paired t-test: t=6.976, n=7 pairs). **L.** Top, example traces of RVM oIPSCs (blue) and vlPAG oEPSCs (red) before and after bath application of DAMGO (1 μM; black). Bottom, opioid sensitivity reported as % suppression of amplitude by DAMGO (RVM oIPSC = 47.6 ± 7.0%, PAG oEPSC = 11.4 ± 4.2%; two-sided unpaired t-test: t=3.785, n=8 RVM IPSC, n=5 PAG EPSC). All summary data reported as mean ± SEM.

To gain insight into the synapses driving these changes in action potential firing, we measured optically-evoked postsynaptic currents in voltage clamp recordings. To electrically isolate excitatory and inhibitory postsynaptic currents (oEPSCs and oIPSCs), we applied light while holding neurons at both –70 mV and 0 mV, respectively. Upon stimulating vlPAG axons (1 x 5 ms pulse), we detected oEPSCs in 30/36 neurons. Seven of these 30 neurons also responded with oIPSCs, but we did not observe any neurons displaying oIPSCs only. The relative oEPSC and oIPSC peak amplitudes are consistent with the overall excitatory drive from vlPAG observed in current clamp (**Figure 5E**). On the other hand, upon stimulating RVM axons, we found oEPSCs in 32/72 neurons and oIPSCs in 62/72 neurons, while 30/72 LC neurons exhibited both. The relative oEPSC and oIPSC peak amplitudes are consistent with the overall net inhibition of LC firing by the RVM (**Figure 5F**).

We next used pharmacology to dissect the underlying synaptic receptors, assess the presence of mono-and/or poly-synaptic connections, and evaluate the mu opioid-sensitivity of each pathway. Consistent with a glutamatergic, monosynaptic excitatory connection, vlPAG-driven oEPSCs were blocked by the AMPA receptor antagonist NBQX (10 µM) and were completely abolished by the voltage-gated sodium channel blocker TTX (1 µM), but could subsequently be rescued by application of the voltage-gated potassium channel antagonist 4-aminopyridine (4-AP; 100 μM) (**Figure 5G-H**). Indicative of a mixed GABAergic and glycinergic projection, RVM-driven oIPSCs were partially blocked by the GABA-A receptor antagonist GABAzine (20 μM), and fully blocked by subsequent addition of the glycine receptor antagonist strychnine (10 μM) (**Figure 5I**). Demonstrating the presence of a monosynaptic inhibitory connection, the oIPSCs were abolished and restored by TTX and 4-AP, respectively (**Figure 5J**). However, bath application of NBQX and the NMDA antagonist CPP (10 µM each) significantly decreased the oIPSC amplitude, suggesting an additional feed-forward, polysynaptic inhibitory component (**Figure 5K**).

Finally, we assayed the opioid sensitivity of vlPAG-driven oEPSCs and RVM-driven oIPSCs by bath applying DAMGO (1 µM) (**Figure 5L**). Whereas vlPAG-driven oEPSCs were only slightly suppressed, RVM-driven oIPSCs were strongly attenuated. This suggests that in the presence of opioids, vlPAG excitatory drive to the LC remains largely intact, whereas suppression of inhibitory synaptic transmission from the RVM is poised to disinhibit LC neurons.

### vlPAG and RVM inputs to LC modulate nociception

We next aimed to determine the contributions of these synaptic pathways to nociception and systemic morphine antinociception. Due to the prominence of branching axons, we used a chemogenetic loss-of-function strategy that restricts inhibition to synaptic terminals in target structures via CNO infusion through implanted cannulas (*62*). We first investigated the contribution of vlPAG output to the LC. After bilateral injection of AAV-DIO-HM4D-mCherry into the vlPAG of *VGLUT2-cre* mice and 3 weeks of expression, we implanted injected mice and control uninjected wildtype mice with cannulas bilaterally over the LC. Following recovery, mice were tested on the hot plate after bilateral infusion of either saline or CNO (3 µM), both in an opioid-naïve state and after injection of morphine (5 mg/kg, *s.c.*). In wildtype mice, intra-LC infusion of CNO had no effect either in the absence or presence of morphine (**Figure 6A**). In vlPAG*^VGLUT2-cre^*::HM4D mice, although intra-LC infusion of CNO did not alter baseline nociception, it produced a large, partial reduction in systemic morphine antinociception (**Figure 6B**). These results indicate that vlPAG glutamatergic output to LC is crucial for systemic morphine antinociception but may work in tandem with other descending structures (*e.g.*, the RVM) to achieve the full morphine effect.

**Figure 6.**
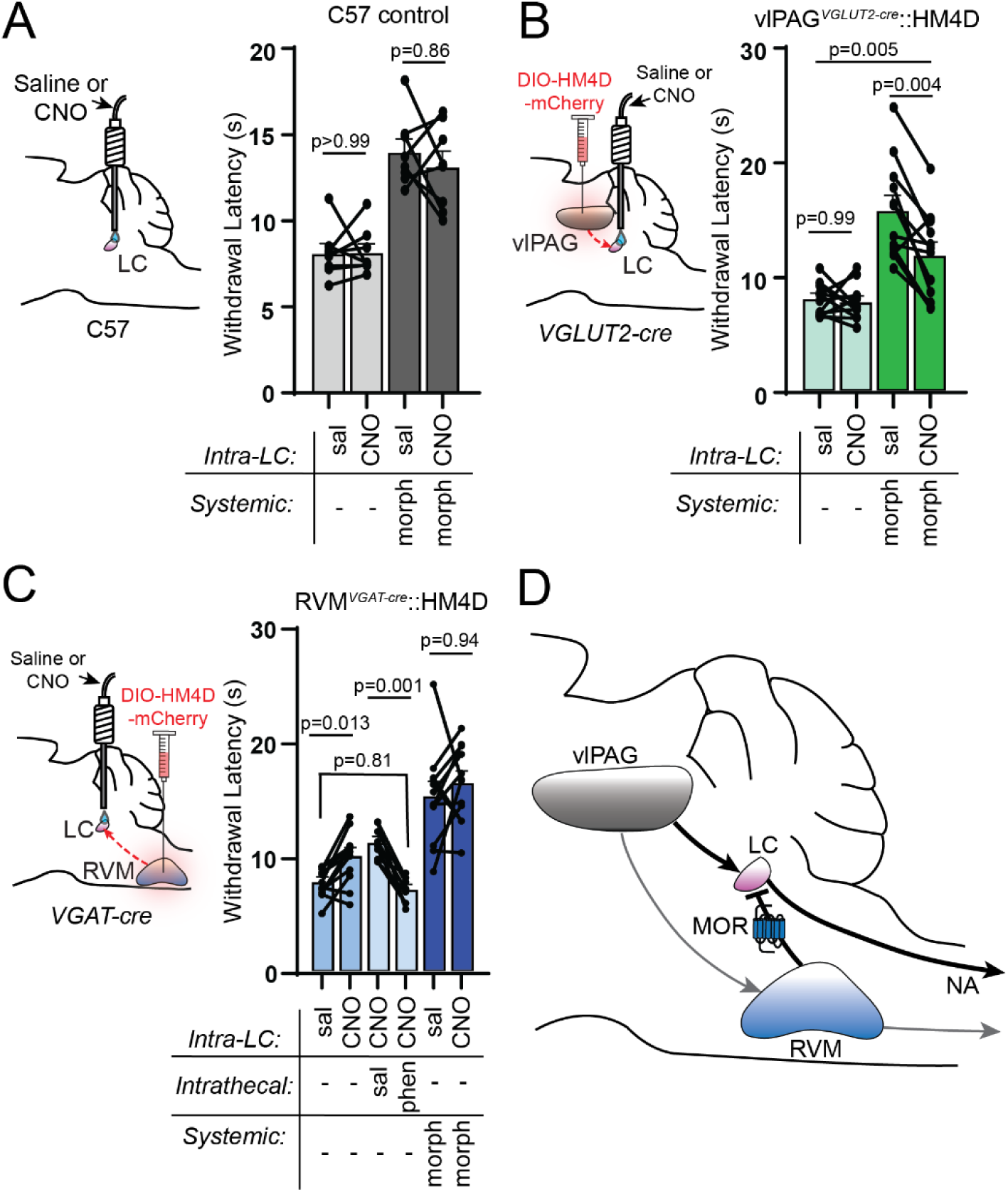
Pathway specific modulation of vlPAG and RVM terminals in LC modulates nociceptive behavior. **A**. Left: cannula placement over bilateral LC of uninjected C57 control mice. Right: hot plate withdrawal latencies of control mice microinfused in LC with saline (150nl) vs. CNO (3 μM 150nl) without (light grey) and with 5 mg/kg morphine, s.c. (dark grey; Two-way repeated measures ANOVA with post hoc Tukey’s multiple comparisons test; n=8 mice; sal vs. CNO effect, p=0.4178, F(1,7)=0.7412, morphine effect, p=0.0001, F(1,7)=57.98; morphine x CNO interaction, p=0.58, F(1,7)=0.3434). **B.** Left: viral injection of AAV-DIO-HM4Di-mCherry in bilateral vlPAG of *VGLUT2-cre* mice with cannula placement over bilateral LC. Right: hot plate withdrawal latencies after microinfusion with saline vs. CNO without (light green) and with 5m/kg morphine, s.c. (dark green; Two-way repeated measures ANOVA with post hoc Sidak’s multiple comparisons test; n=12 mice; sal vs. CNO effect, p=0.0022, F(1,11)=15.72, morphine effect, p=0.0003, F(1,11)=27.60; morphine x CNO interaction, p=0.0106, F(1,11)=9.456). **C.** Left: viral injection of AAV-DIO-HM4Di-mCherry in bilateral RVM of *VGAT-cre* mice with cannula placement over bilateral LC. Right: hot plate withdrawal latencies after microinfusion of saline vs. CNO (blue bars n=12 mice), microinfusion of CNO with intrathecal injections of saline vs. phentolamine (5ug, light blue bars, n=9 mice), and microinfusion of saline vs. CNO with 5 mg/kg morphine, s.c. (dark blue, n=12 mice; Mixed effects analysis with matching across row and post hoc Tukey’s multiple comparisons test, p<0.0001, F(2.560,25.09)=27.36). **D.** Circuit diagram of DPMS inputs to LC and their opioid sensitivity. Data in each graph reported as mean ± SEM.

We next sought to determine if RVM-to-LC inhibitory projections shape nociception. We first challenged RVM*^VGAT-cre^*::HM4D mice with systemic CNO to determine the effect of inhibiting all RVM inhibitory neurons. This global inhibition had no effect on baseline nociception or morphine (5 mg/kg) antinociception on the hot plate (**Figure S5**). In contrast, suppression of RVM inhibitory output to the LC via local CNO infusion increased hot plate withdrawal latencies, which is consistent with disinhibition of descending LC-NA neurons (**Figure 6C**).

Confirming this hypothesis, intrathecal phentolamine completely blocked the resulting antinociception. Chemogenetic suppression of this pathway had no effect on morphine antinociception, which is consistent with occlusion of MOR-mediated synaptic suppression by HM4D activation (and vice-versa). Taken together, these pathway-specific manipulations are consistent with direct excitation of LC by vlPAG glutamatergic neurons and tonic inhibition by RVM inhibitory neurons that leads to opioid-driven disinhibition in the presence of opioids to shape nociceptive behavior (**Figure 6D**).

## Discussion

In this study, we establish a critical and previously underappreciated role for LC activity in systemic morphine antinociception and delineate the synaptic mechanisms by which vlPAG and RVM activate the LC in response to supraspinal opioid signaling. Although the LC is frequently included in DPMS circuit models, prior lesion studies aimed at identifying the sources of spinal NA concluded that the LC makes only a minor contribution, which led to a focus on the A7 nucleus (*28*, *63–66*). In contrast, our results establish a central role for the LC in morphine antinociception in mice. We find that systemic morphine induces c-Fos in LC NA neurons, consistent with neuronal activation, although other morphine-responsive receptors, such as Gs-coupled Mas-related G protein-coupled receptors (MRGs) (*67–69*) could be involved. However, this effect was also significantly blocked by systemic opioid antagonism, pointing to a role for upstream opioid receptors in vlPAG interneurons and/or the inhibitory RVM inputs described in this study. Importantly, our loss-of-function manipulations indicate a major contribution of the LC to systemic morphine antinociception. The apparent discrepancy between our findings and previous work might be attributed to differences in the anatomy and/or neurochemistry of the DPMS in rats and mice. Indeed, in our hands, mouse LC neurons exhibited a comparatively small opioid response, whereas the rat LC is profoundly inhibited by opioids (*42*, *43*). In either case, because transgenic mice are widely used in contemporary pain research, understanding their descending pain modulatory circuitry is of great importance.

The vlPAG has been long appreciated to contribute to morphine antinociception due to the initial finding that local vlPAG opioid administration produces potent antinociception. However, its role has not been unequivocally established, as lesion studies in rats arrive at different conclusions (*70*, *71*), perhaps due to differences in lesion protocol and the limited precision of this method.

Using cell type-specific chemogenetic loss-of-function, our work demonstrates that glutamatergic output from the vlPAG is critical for morphine antinociception. This is consistent with classic models of opioid-mediated disinhibition of vlPAG neurons that project to the RVM, which we found to send prominent axon collaterals to the LC. This surprising finding suggests that activation of antinociceptive projections neurons in the vlPAG recruits the LC and RVM in parallel. This model is further bolstered by the relatively low mu opioid-sensitivity of vlPAG excitatory synaptic transmission onto LC neurons, such that disinhibited vlPAG output should faithfully drive LC activity in the presence of systemic MOR agonists.

Our results also reveal several distinctions between the role of NA in spinal and supraspinal opioid antinociception. At low doses of morphine (*e.g.*, 5 mg/kg), the antinociception is mediated primarily by supraspinal opioid receptors, as evidenced by nearly complete loss of systemic morphine antinociception upon chemogenetic vlPAG silencing and loss of descending NA. In contrast, higher morphine doses (>10 mg/kg) may directly engage spinal opioid receptors, as intrathecal phentolamine only partially blocks this antinociception. These results are consistent with a proposed multiplicative effect of supraspinal and spinal opioids at lower morphine doses, whereas the effect of either site can mediate antinociception at higher doses (*39*, *72*, *73*). Yet, within the spinal cord, NA-opioid interactions are likely critical. In our data, both vlPAG*^VGLUT2-Cre^*::HM3D and low-dose systemic morphine antinociception were completely blocked by phentolamine *or* naltrexone. Since both involve the vlPAG, one possibility is that spinal endogenous opioids are recruited via the RVM. Alternatively, because the antinociception produced by intrathecal NA is blocked by intrathecal opioid antagonists (*74*), NA may drive endogenous opioid release in the spinal cord. However, neither spinal NA nor spinal opioids alone are sufficient to support the vlPAG-driven antinociception. Intrathecal naltrexone blocked systemic morphine antinociception at all doses tested, consistent with spinal opioid signaling being a critical end-point. Because intrathecal morphine antinociception does not require spinal NA signaling, we posit that spinal opioid receptors gate spinal NA-mediated antinociception.

Our results indicate a major role for the LC, as well as the glutamatergic vlPAG projections to it, in systemic morphine antinociception. However, upon inhibition of either, abrogation of antinociception was incomplete. This suggests that other parallel descending pathways also contribute to systemic morphine effects. A likely possibility is that vlPAG projections to RVM contribute to morphine antinociception in a parallel and additive fashion, taking advantage of vlPAG bifurcations to both structures. Alternatively, A5 and A7 are other sources of descending NA that may be involved in morphine antinociception. A large body of literature supports an antinociceptive function for A7 that depends on spinal NA signaling. More recently, A5 has been proposed to mediate diffuse noxious inhibitory control of pain downstream of the LC (*75*, *76*).

Both of these regions also receive anatomical inputs from both vlPAG and RVM (*29*, *34*). Future experiments are required to assess the contributions of these pathways to systemic opioid antinociception.

Finally, our discovery of a largely inhibitory drive from RVM to LC was unexpected. While chemogenetic inhibition of these RVM outputs specifically to LC caused spinal NA-dependent antinociception, presumably via disinhibition of LC, inhibiting all RVM inhibitory neurons had no effect on either opioid-naïve nociception or morphine antinociception (**Figure S5**). This may be due to interactions between pro– and antinociceptive RVM projections to the spinal cord (*2*, *35*). Interestingly, LC-projecting RVM neurons do not appear to project to the spinal cord and instead send ascending projections to the thalamus. It is not currently clear how LC-projecting RVM neurons might fit into the canonical On– and Off-cell framework. Additionally, how these LC-projecting RVM neurons become activated remains to be determined.

Overall, this work establishes the LC as a central source of spinal NA that generates systemic morphine antinociception and identifies excitatory input from vlPAG neurons as the critical synapse that drives LC in this context. In addition, it establishes a new synaptic pathway between the medial RVM and LC that can produce acute antinociception via disinhibition of LC-NA neurons, leading to spinal NA release. Although this study utilized an exogenous opioid drug, the circuit elements uncovered are likely involved in other forms of top-down descending pain modulation that rely on endogenous opioids, such as placebo– and stress-induced analgesia.

## Materials and Methods

### Animals

All procedures were performed in accordance with protocols approved by the University of California San Diego Institutional Animal Care and Use Committee (UCSD IACUC protocol S16171) and guidelines from the US National Institutes of Health Guide for Care and Use of Laboratory Animals. Mice were group housed, maintained on a 12-hour reversed light/dark cycle, and allowed *ad libitum* access to food and water. Experiments were performed under red lighting during the dark period. Strains used include the following: C57Bl/6J (Jackson Laboratory stock #664), *DBH*-cre (MMRRC/GENSAT #032081-UCD, Tg(DBH-cre)HK212Gsat/Mmucd), *VGLUT2-*IRES-cre (Jackson Laboratory stock #028863, B6J.129S6(FVB)-Slc17a6^tm2(cre)Lowl^/MwarJ), and *VGAT*-IRES-cre (Jackson Laboratory stock #028862, B6J.129S6(FVB)-Slc32a1^tm2(cre)Lowl^/MwarJ). Ai14 tdTomato (Jackson Laboratory stock #7914, B6.Cg-Gt(ROSA)^26Sortm14(CAG-tdTomato)Hze^/J) and Oprm1^fl/fl^ (Jackson Laboratory stock #30074, B6;129-Oprm1^tm1.1Cgrf^/KffJ) mice were also obtained from Jackson Labs, but bred in-house to *DBH*-cre mice. Mice were used for experiments between the ages of 8-20 weeks. Both male and female mice were used for all experiments.

### Drugs

The following drugs were purchased from HelloBio: CNO (HB1807), NBQX disodium salt (HB0443), Tetrodotoxin citrate (HB1035), GABAzine (SR 95531 hydrobromide; HB0901), and (R)-CPP (HB0021). Naltrexone hydrochloride (N3136), naloxone hydrochloride (N7758), 4-aminopyridine (A78403) and strychnine hydrochloride (S8753) were purchased from Sigma-Aldrich. Morphine sulfate was purchased from Spectrum Chemicals (M1167). Phentolamine hydrochloride was purchased from Abcam (ab120791). DAMGO was purchased from R&D Systems, Inc. (1171). CNV-Y-DAMGO was prepared in house, as previously reported (*44*). Drugs to be injected or intracranially infused were dissolved in 0.9% saline and sterile filtered before use.

### Viral constructs

The following viruses were purchased from Addgene: AAV8-hsyn-DIO-HM4Di-mCherry, (Addgene 44362, titer 2.3×10^13^), AAV8-hsyn-DIO-HM3Dq-mCherry (Addgene 44361, titer 2.1×10^13^), AAVretro-hsyn-cre (Addgene 105553, titer 2.1×10^13^), AAVretro-Ef1a-FlpO (Addgene 55637, titer 1.3×10^13^). The following were made and titered in-house using Addgene plasmids: AAVDJ-hsyn-DIO-mCherry (Addgene 50459, titer 3.5×10^12^), AAVDJ-ef1a-fDIO-EYFP (Addgene 55641, titer 2×10^11^), AAVDJ-Ef1a-mCherry-IRES-cre (Addgene 55632, titer 4.3×10^12^), AAVDJ-ef1a-DIO-ChR2(H134R)-mCherry (Addgene 20297, titer 2.55×10^13^). AAV1-FLEX-ef1a-taCasp3-TEVP (titer 2.1×10^12^) and AAV1-CAG-FLEX-tdTomato (titer 7.6×10^12^) were both purchased from the UNC Vector Core. We received AAVDJ-CAG-DIO-Kir2.1-P2A-zsGreen as a gift from the laboratory of Dr. Byungkook Lim.

### Intrathecal injections

Intrathecal injections were performed according to the protocol detailed by Hylden and Wilcox (*77*). Acute percutaneous intrathecal injections were executed using a 30G 1-inch needle attached to a 10 ul Hamilton syringe via polyethylene tubing. Mice were lightly anesthetized with 2% isoflurane, the fur over their lumbar spine was removed with electric clippers, and the skin disinfected with alternating povidone-iodine solution and isopropyl alcohol. While firmly holding the pelvic girdle, the needle was inserted into the skin over the lumbar spine at a 20-degree angle. The needle was guided between the vertebrae until it entered the spinal column, which was signified by a tail flick. Each intrathecal injection was given at a volume of 5 μl. Mice were given at least 10-15 minutes to recover from the intrathecal injection before behavioral testing.

### Behavior assays

#### Thermal nociceptive behavior

Mice were habituated to a hot plate (Thermo Fisher Scientific #SP88857100) at room temperature in a clear plastic cylinder for at least 20 minutes. During testing, mice were placed on the hot plate at 52°C and observed for a nocifensive response (hind paw withdrawal, shaking, licking, or jumping). Mice were immediately removed from the hot plate at the first sign of a nocifensive response, and the time to paw withdrawal was recorded. Each trial was terminated after a maximum of 60 seconds, even if no withdrawal occurred, to avoid tissue damage. Mice were tested twice on the hot plate during each session with a 3-5-minute intertrial interval.

#### Mechanical nociceptive behavior

Mice were tested for mechanical thresholds using either the von Frey or pin prick assay. In both cases, mice were placed in a clear cylinder on an elevated wire grid and allowed to habituate for 20 minutes. Using the up-down method (*78*), von Frey filaments (Ugo Basile) of varying stiffnesses were applied to each hindpaw separately, with a 3-minute break until returning to the first paw, until a withdrawal response could be recorded and a 50% withdrawal threshold could be calculated. The resulting threshold values (in grams) for each paw were averaged together to calculate the mechanical threshold for each mouse. During the pin prick assay, a fine insect pin (size 000, ThermoFisher Scientific NC9295307) was applied to the plantar surface of each hind paw 5 times at 5-minute intervals for a total of 10 trials. Response to pin was calculated as the number out of 10 trials in which the mouse displayed nocifensive withdrawal behavior.

#### Elevated plus maze and open field

Mice were placed on an elevated plus maze with two open and two closed arms (52 cm diameter) and video recorded (Logitech) for a 20-minute period. The SMARTv3.0 video tracking software (Panlab) was used to measure the percent of time spent in the open vs. closed arms of the apparatus. Additionally, the distance traveled during the 20-minute testing period was reported. Similarly, locomotion in the open field was tested by placing mice in a square arena (18 cm x 18 cm) for a 20-minute testing period and the SMARTv3.0 software was used to track mouse trajectory and calculate distance traveled.

### Surgeries

Before surgery, mice were deeply anesthetized by induction at 5% isoflurane, after which anesthesia was maintained by 2% isoflurane (SomnoSuite, Kent Scientific). After mice were placed in a stereotaxic frame (David Kopf Instruments), a midline incision was made through the scalp following fur removal and site preparation by alternating povidone-iodine and 70% isopropyl alcohol. 100-250 nl of virus was injected at a rate of 100 nl/min at defined stereotaxic coordinates. The stereotaxic coordinates used for viral injections are as follows: vlPAG, angle ±10°, AP –4.60 mm, ML ± 0.32 mm, DV 2.85 mm; RVM, angle 10°, AP –7.00 mm, ML +0.26 mm and –0.67 mm, DV 6.30 mm and 6.26 mm; LC, angle ±10°, AP –5.45, ML ±0.75 mm, DV 4.10 mm. For bilateral LC cannula implants, the following coordinates were used: angle ±15°, AP –5.45 mm, ML ±0.9 mm, DV 3.80 mm. Guide cannulas were 26G, included a 5 mm pedestal, and were cut 5 mm below the pedestal. Internal cannulas were 33G and included 1 mm projection from the end of the guide cannula (Plastics One/Protech International, Inc.). Cannulas were secured to the surface of the skull using light-cured dental epoxy and anchored by one screw. For all surgeries, mice were administered 5 mg/kg ketoprofen (MWI Veterinary Supply) before the end of surgery and 24 hours later and monitored for recovery for 5 days.

### Brain slice preparation

Mice were anesthetized with isoflurane before rapid decapitation. Brains were removed, blocked, and mounted in a VT1000s vibratome (Leica Instruments). Coronal midbrain slices (190 μm) containing the LC were prepared in ice-cold choline-based artificial cerebrospinal fluid containing 25 mM NaHCO_3_, 1.25 mM NaH_2_PO_4_, 2.5 mM KCl, 7 mM MgCl_2_, 25 mM glucose, 0.5 mM CaCl_2_, 110 mM choline chloride, 11.6 mM ascorbic acid, and 3.1 mM pyruvic acid, equilibrated with 95% O_2_/5% CO_2_. Slice were transferred to 32°C oxygenated artificial cerebrospinal fluid (ACSF) containing 125mM NaCl, 2.5 mM KCl, 25 mM NaHCO_3_, 1.25 mM NaH_2_PO_4_, 2 mM CaCl_2_, 1 mM MgCl_2_, and 10 mM glucose, osmolarity 290. Slices were incubated for 20 minutes, then brought to room temperature before recording.

### Slice Electrophysiology

Slice physiology experiments were performed in a chamber continuously perfused with warmed (32°C) ACSF equilibrated with 95% O_2_/5% CO_2_. Recording pipettes were pulled from borosilicate glass on a P-1000 Flaming/Brown micropipette puller (Sutter Instruments) to a resistance of 1-3.5 MΩ. Recordings were made with an Axopatch 700B amplifier (Axon Instruments) and data sampled at 10 kHz, filtered at 3 kHz, and acquired using National Instruments acquisition boards and a custom version of ScanImage written in MATLAB (MathWorks). LC neurons were visually identified and confirmed to exhibit tonic action potential spiking of ∼1.5 Hz. Recordings were biased toward the ventral portion of LC on each coronal slice. Recordings of action potential spiking were made in current clamp with patch pipettes filled with an internal solution containing 135 mM KMeSO_3_, 5 mM KCl, 5 mM HEPES, 4 mM MgATP, 0.3 mM NaGTP, 10 mM phosphocreatine, and 1.1 mM EGTA (pH 7.25, 290 mOsm kg^-1^). Firing rate vs. current input (f-I) curves were constructed using 1-second steps of direct current input (–100, 50, 100, 150, 200, 250, 300 pA) 10 seconds apart. The effect of CNV-Y-DAMGO circulating in the bath on spiking was determined by presenting a single 50 ms flash of UV light (365 nm LED, 84 mW, pE-300ultra, CoolLED) and recording tonic spiking for 20 seconds before and 100 seconds after the flash. Firing rate in the 10 seconds before and after the flash, latency to spike after the flash as a measure of the pause in action potential firing, and the change in membrane potential after the flash were measured. For the effect of optogenetically-mediated excitation of inputs to LC on tonic spiking, 50×2 ms pulses of blue light (470 nm, 18 mW, pE-300ultra LED) at 25 Hz for a total of 2 seconds were delivered. The firing rate in the 2 seconds before and 2 seconds during blue light stimulation were used to compare light-evoked excitation or inhibition.

Recordings of evoked postsynaptic currents were made in voltage clamp with patch pipettes filled with an internal solution containing 135 mM CsMeSO_3_, 3.3 mM QX314 Cl^-^ salt, 10 mM HEPES, 4 mM MgATP, 0.3 mM NaGTP, 8 mM phosphocreatine, and 1 mM EGTA (pH 7.2-7.3, 295 mOsm kg^-1^). Cells were rejected if holding currents became more negative than –200pA or if series resistance exceeded 25 MΩ. Recordings of opioid-evoked currents were recorded in the presence of NBQX (10 μM), CPP (10 μM), GABAzine (20 μM), and strychnine (10 μM) to eliminate synaptic currents. As in current clamp, CNV-Y-DAMGO was photoactivated by a single 50 ms flash of UV light. While recording optogenetically-evoked EPSCs and IPSCs in LC neurons, ChR2 terminals were stimulated with a single 5 ms flash of blue light. Initial characterization of oEPSCs and oIPSCs was done in the absence of synaptic blockers and each current was isolated by holding the LC neuron at –70 mV or 0 mV, respectively. Further pharmacological verification of evoked currents was performed with AMPA blocker NBQX (10 μM), NMDA blocker CPP (10μM), GABA-A receptor blocker GABAzine (SR 95531, 20μM), glycine receptor blocker strychnine (10 μM), sodium channel blocker TTX (1 μM), and potassium channel blocker 4-aminopyridine (100 μM). Opioid sensitivity of evoked postsynaptic currents was assessed by bath perfusion of mu-opioid receptor agonist DAMGO (1 μM). All electrophysiology data were processed in Igor Pro (Wavemetrics).

### Fluorescent *in situ* hybridization (FISH)

FISH was performed either in naïve wild type mice or in mice bilaterally injected in RVM with green retrobeads (Lumafluor, Inc.). Mice were deeply anesthetized and decapitated. Brains were quickly removed and frozen in Tissue-Tek OCT medium (Sakura) on dry ice until completely solid. Brain slices (8 μm) were prepared on a cryostat (Leica CM 1950) and adhered to SuperFrost Plus slides (VWR). Samples were fixed with 4% paraformaldehyde and processed according to instructions in the ACD Bio RNAscope Fluorescent Multiplex Assay (Fluorescent Reagent Kit v2) manual and cover slipped with ProLong antifade media (Molecular Probes).

Images were taken on a Keyence microscope (BZ-X710) using a 60× 1.4 NA oil immersion objective configured for structured illumination microscopy. Puncta counting in FISH images was completed using a custom pipeline designed in CellProfiler.

### Histology

For all other mice, brains were fixed using chilled 4% paraformaldehyde in phosphate buffered saline (PBS) by transcardial perfusion and cryoprotected in 30% sucrose in PBS solution. In some cases, spinal cords were also removed and processed in the same fashion. Brain slices (40 μm) were prepared on a freezing microtome (ThermoFisher Scientific Microm HM450). For sections selected for immunohistochemistry, slices were blocked in PBS-Triton (0.3% TritonX-100 in 1× PBS) with 5% donkey or goat serum for 1 hour at room temperature. Incubation in primary antibodies occurred in PBS-Triton plus 1% serum at 4°C for 24-72 hours. Following 3 10-minute wash steps in PBS, the slices were incubated in PBS-Triton plus 1% serum with secondary antibodies for 4 hours at room temperature. After 3 more wash steps, slices were mounted and cover slipped with mounting media containing DAPI (Vector Laboratories, H1200). Primary antibodies (1:500) used include rabbit anti-c-Fos (9F6) (#2250, Cell Signaling Technology), rabbit anti-tyrosine hydroxlase (AB152, EMD Millipore), and mouse anti-tyrosine hydroxylase (T2928, Sigma). Secondary antibodies used include Alexa Fluor 488 Donkey anti-rabbit, Alexa Fluor 555 donkey anti-mouse, Alexa Fluor 488 goat anti-rabbit and Alexa Fluor 555 goat anti-rabbit (1:500, Invitrogen). After mounting, slices were imaged on a Keyence microscope (BZ-X710) at 2×, 10×, or 20× magnification. Cell counts and quantification of fluorescent colocalization were performed using custom pipelines in Cell Profiler. Pixel intensity analysis was performed in Image J, z-scores calculated in Microsoft Excel, and resulting values graphed in GraphPad Prism.

## Statistics

Data in each figure are presented as mean ± SEM and individual data points from experimental replicates are plotted in most cases. Statistical analyses were performed in GraphPad Prism. The test performed, number of experimental replicates, mean ± SEM for each condition, statistics provided by ANOVA and two-way ANOVA tests for each variable, and p-values for the overall test are provided in the figure legend. Individual p-values resulting from post hoc tests to correct for multiple comparisons appear in the figures. Repeated measures within individual replicates were taken into account where applicable based on experimental design. Data were tested for normality using the Shapiro Wilk test, and non-parametric statistics were used when necessary and possible. In certain cases, when multiple conditions included different numbers of experimental replicates, a mixed-effects model was used in place of a two-way ANOVA. Raw data for each graph as well as the results of statistical tests are available in the Source Data spreadsheet.

## Acknowledgements

We thank B.K. Lim for providing the Kir2.1 virus, A.E. Layden & J. Chang-Weinberg for genotyping support, and B.K. Lim, S. Han, and W. Campana for helpful discussions.

## Author Contributions

STL, GL, TLY, and MRB conceived the project and designed the work. STL, JP, and JCY performed experiments and analyzed data. STL, TLY, and MRB wrote and edited the manuscript.

## Data and Materials availability

Correspondence and material requests should be addressed to Dr. Matthew Banghart. All data, including complete statistics tables, are available in the supplementary materials Source Data file.

## Supplementary Materials

**Figure S1.**
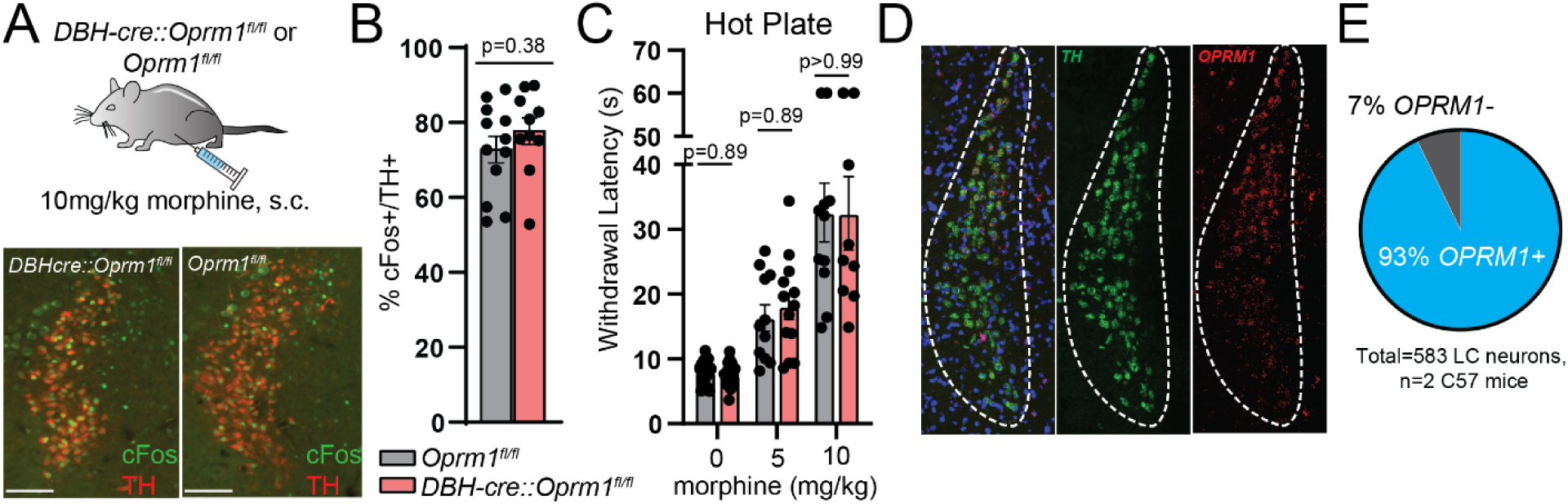
MORs in LC are not responsible for morphine induced activity changes or behavior. **A**. Top: 10 mg/kg, s.c. morphine injections in *DBH-cre::Oprm1^fl/fl^* mice and control *Oprm1^fl/fl^*littermates. Bottom: Representative images of morphine-induced c-Fos expression (green) and immunhistochemical TH LC labeling (red). **B.** Percentage of TH+ LC neurons that colocalize with green c-Fos signal (5-8 images analyzed per mouse; n=10 *DBH-cre::Oprm1^fl/fl^*, n=12 *Oprm1^fl/fl^*; *DBH-cre::Oprm1^fl/fl^* = 77.6 ± 3.5%, *Oprm1^fl/fl^* = 72.7 ± 3.5%; p=0.38, two-sided Mann-Whitney test). **C.** Hot plate withdrawal latencies compared between *DBH-cre::Oprm1^fl/fl^* and *Oprm1^fl/fl^* littermates at 0, 5, and 10 mg/kg doses of morphine, s.c. (Mixed effects analysis with post hoc Sidak’s multiple comparisons test; baseline n=18 *DBH-cre::Oprm1^fl/fl^*, n=18 *Oprm1^fl/fl^*; 5 mg/kg morphine n=13 *DBH-cre::Oprm1^fl/fl^*, n=11 *Oprm1^fl/fl^*; 10 mg/kg morphine n=9 *DBH-cre::Oprm1^fl/fl^*, n=11 *Oprm1^fl/fl^*; morphine dose effect, p<0.0001, F(1.261, 25.22)=50.93; genotype effect, p=0.835, F(1,34)=0.044; morphine dose x genotype interaction, p=0.88, F(2,40)=0.1283). **D.** Representative images of fluorescent *in situ* hybridization of LC with probes against *TH* (green) and *OPRM1* (red). **E.** Quantification of the percentage of *TH*+ LC neurons that express *OPRM1* transcripts.

**Figure S2.**
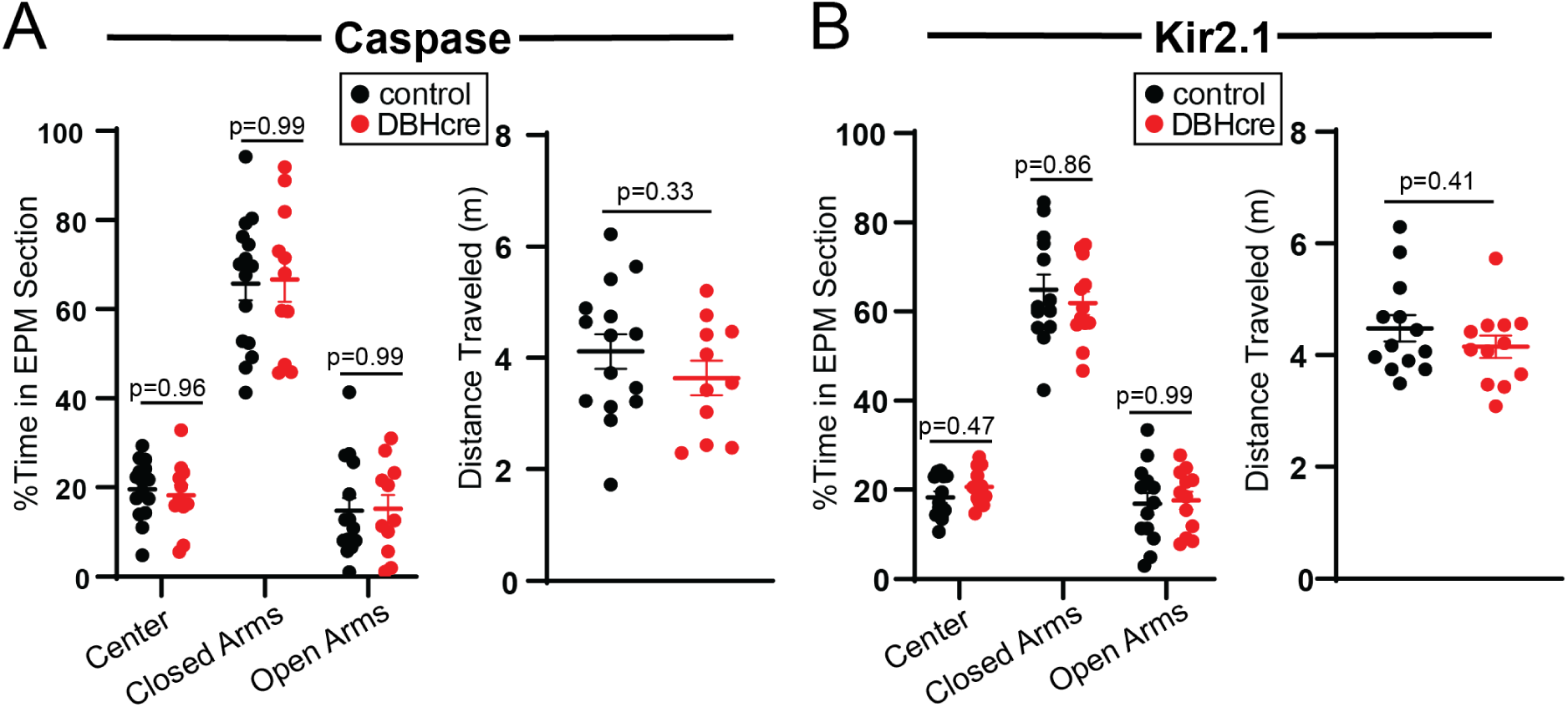
LC ablation or silencing does not affect baseline anxiety behavior or locomotion on the elevated plus maze. **A**. Left: percentage of time spent in different sections of the elevated plus maze (EPM) during a 20 minute session in *DBH-cre*::tdTom and control *TdTom* mice injected with AAV-DIO-Casp3 (same cohort that underwent hot plate and pin prick behavior in Figure 2F; Two-way repeated measures ANOVA with post hoc Sidak’s multiple comparisons test; n=11 *DBH-cre* mice, n=15 control; genotype effect, p=0.489, F(1,24)=0.4934). Right: distance traveled during the 20-minute EPM session (two-sided Mann-Whitney test). **B.** Same as in A, but for AAV-DIO-Kir2.1-zsGreen-injected mice from Figure 2J-K. Left: Two-way repeated measures ANOVA with post hoc Sidak’s multiple comparisons test; n=12 *DBH-cre* mice, n=13 control; genotype effect, p=0.386, F(1,23)=0.7797. Right: two-sided Mann-Whitney test.

**Figure S3.**
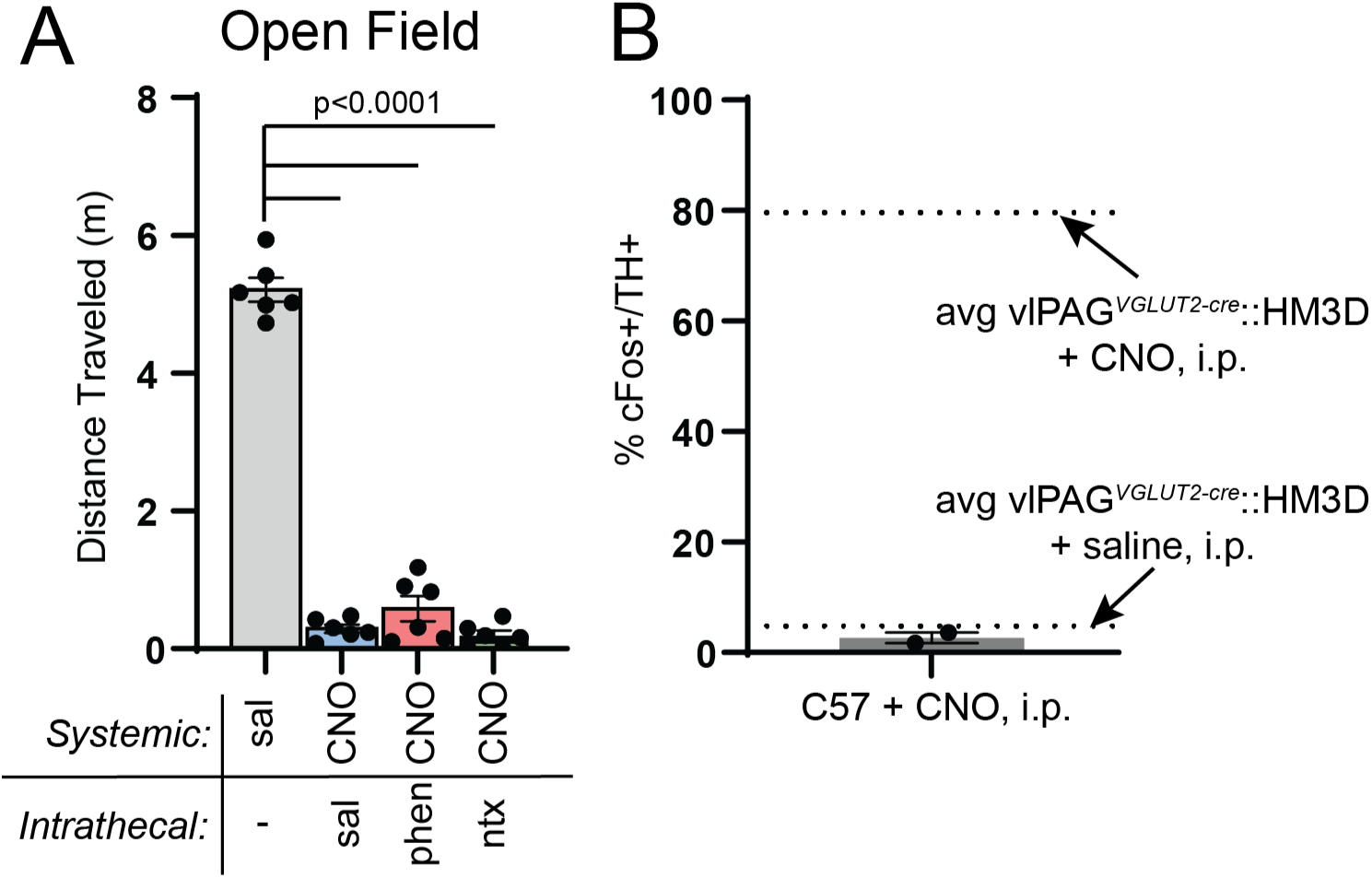
Chemogenetic PAG activation leads to decreased locomotion that can be decoupled from antinociception. **A**. Distance traveled (m) during a 30-minute open field session after systemic injection of saline or CNO (3 mg/kg, i.p.) with intrathecal saline, phentolamine (5 μg), or naltrexone (5 μg) in vlPAG^VGLUT2-cre^::HM3D mice (Repeated measures one-way ANOVA with post hoc Tukey’s multiple comparisons test, p<0.0001, F(2.467, 12.33)=392.1). **B.** LC c-Fos expression induced by 3 mg/kg, i.p. CNO injection in uninjected wild type mice expressed as % TH+ LC neurons that colocalize with c-Fos. Dashed lines represent the average c-Fos induction by CNO and saline in the LC of vlPAG^VGLUT2-cre^::HM3D mice in Figure 3E.

**Figure S4.**
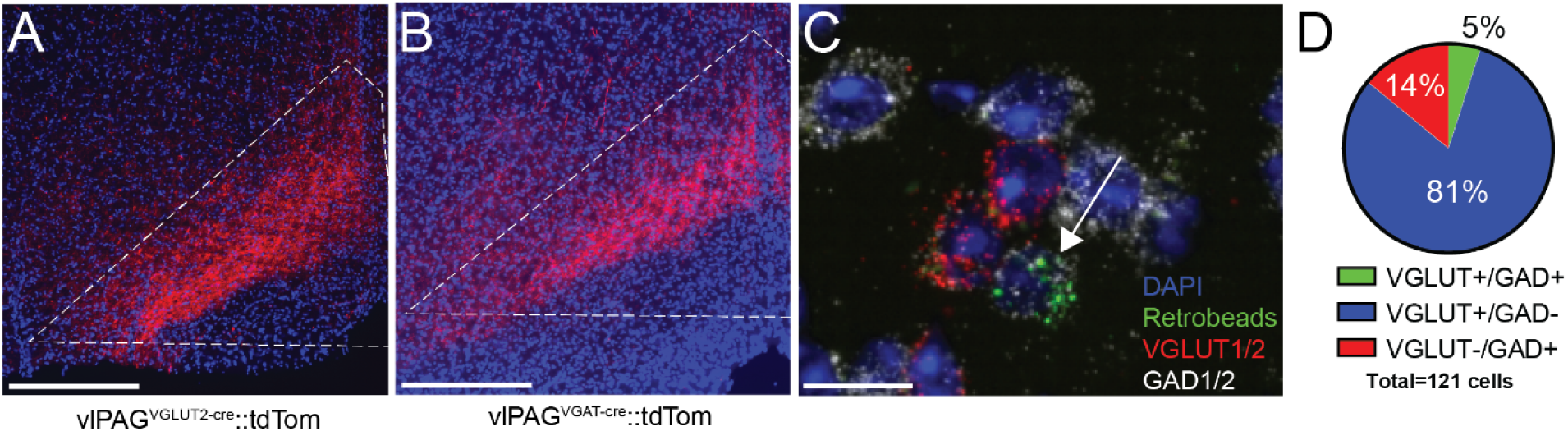
vlPAG sends GABAergic projections to RVM. **A**. Representative image of glutamatergic terminals in RVM (red) of a *VGLUT2-cre* mouse injected in vlPAG with AAV-DIO-tdTom. Scale bar 300 μm. **B.** Representative image of GABAergic/glycinergic terminals in RVM (red) of a *VGAT-cre* mouse injected in vlPAG with AAV-DIO-tdTom. Scale bar 300 μm. **C.** Representative image of results from the retro-FISH method depicting vlPAG with colocalization of retrobeads from RVM (green) and *GAD1/2* transcripts (white) within a single neuron, as well as nearby neurons expressing *VGLUT1/2* transcripts (red). Scale bar 10 μm. **D.** Quantification of *VGLUT1/2* and *GAD1/2* expression within retrobead-labeled vlPAG neurons.

**Figure S5.**
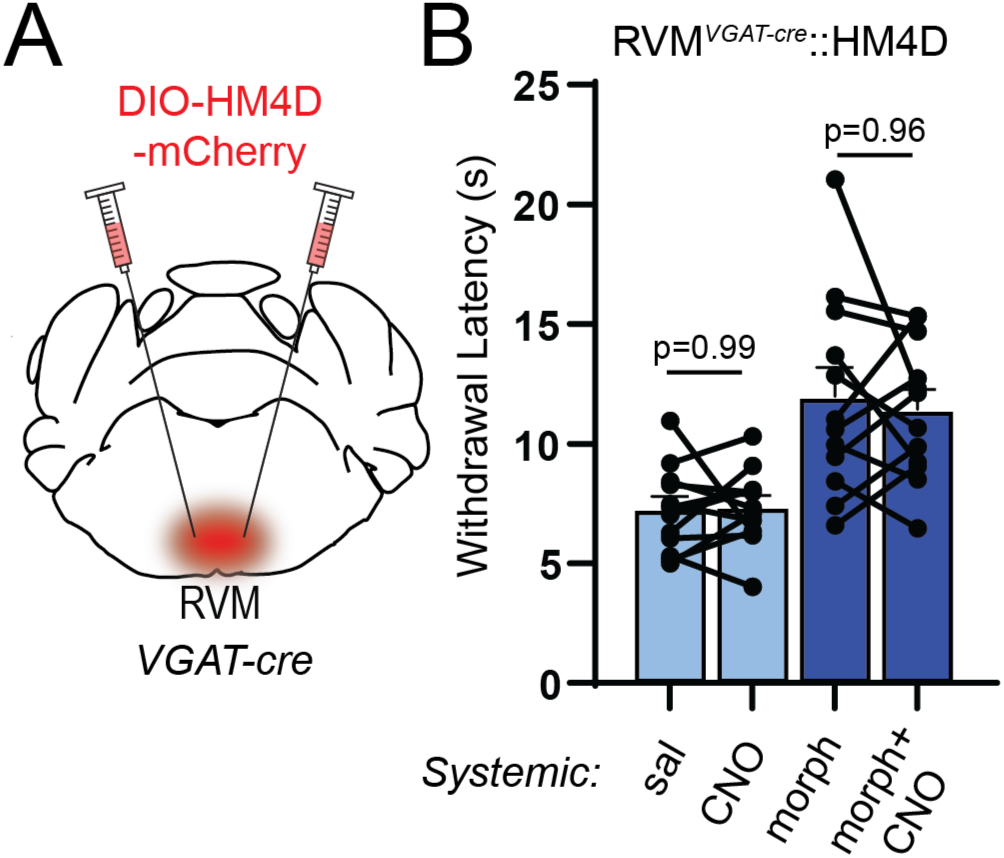
Inhibition of RVM GABAergic neurons does not affect baseline pain behavior or morphine antinociception. **A**. Viral injections of AAV-DIO-HM4Di-mCherry into bilateral RVM of *VGAT-cre* mice. **B.** Hot plate withdrawal latencies of RVM*^VGAT-cre^*::HM4D mice administered 3 mg/kg CNO, i.p. vs. saline without (light blue) and with 5 mg/kg morphine, s.c. (dark blue bars, Two-way repeated measures ANOVA with post hoc Tukey’s multiple comparisons test; n=12 mice; saline vs. CNO effect, p=0.725, F(1,11)=0.1307, morphine effect, p<0.0001, F(1,11)=36.62, morphine x CNO interaction, p=0.60, F(1,11)=0.2897).

